# Snx4-mediated nucleophagy targets transcription factors controlling *ATG* gene expression

**DOI:** 10.1101/2020.05.27.118315

**Authors:** Sara E. Hanley, Stephen D. Willis, Katrina F. Cooper

## Abstract

Autophagy is controlled in part by the repression and activation of Autophagy-related (*ATG*) gene transcription. Here, we demonstrate that the conserved Cdk8 Kinase Module (CKM) of the mediator complex represses transcription of several *ATG* genes. To relieve this repression following nitrogen starvation, Med13 is rapidly degraded via a novel selective autophagy mechanism. This pathway requires the core autophagy machinery but is independent of known nucleophagy systems. It requires the cytosolic filament nucleoporin Gle1, the sorting nexin Snx4-Atg20 heterodimer, and the scaffold protein Atg17. This suggests a model where Med13 traverses through the nuclear pore complex, passing from Gle1 to Snx4. Snx4 then transports Med13 to autophagosomes by binding to Atg17. This previously unidentified nucleophagy pathway also mediates the autophagic degradation of two transcriptional activators of *ATG* genes (Rim15, Msn2) suggesting that this mechanism targets transcription factors that regulate ATG expression. This system provides a new level of selectivity, permitting the cell to fine-tune the autophagic response by controlling the turnover of both positive and negative *ATG* transcription factors.

## INTRODUCTION

Macro-autophagy (hereafter autophagy) is a controlled catabolic process that aids cellular survival during adverse conditions such as starvation or proteolytic stress, by degrading damaged or unnecessary proteins in the vacuole (lysosome in higher eukaryotes) (Klionsky & Codogno, 2013). In budding yeast, non-selective pathways are upregulated in response to nitrogen depletion. This triggers a cascade of events, resulting in non-specific cytosolic cargos being sequestered within autophagosomes and ultimately degraded by vacuolar proteolysis. Selective autophagy pathways use receptor proteins to recognize and deliver specific cargos such as organelles, protein aggregates, and large multi-subunit complexes to autophagosomes (Farre & Subramani, 2016).

Nuclear autophagy or nucleophagy is the least well understood of the selective autophagy mechanisms. Underscoring its importance, various pathologies namely cancer, and neurodegeneration are linked with perturbed nucleophagy (Fu *et al*, 2018). It is best characterized in yeast where macro-nucleophagy utilizes a receptor protein and involves the sequestration of a portion of the nucleus into autophagosomes (Mochida *et al*, 2015). In contrast, micro-nucleophagy (Piecemeal nucleophagy), is autophagosome independent, and forms nuclear-vacuole junctions which pinch off portions of the nucleus directly into the vacuolar lumen (Roberts *et al*, 2003). Recently, an autophagic mechanism has been described that removes defective nuclear pore complexes (NPCs) (Lee *et al*, 2020).

In *S. cerevisiae*, 41 unique autophagy-related (*ATG*) genes have been identified that control this highly coordinated and complex process (Delorme-Axford & Klionsky, 2018). Accordingly, these genes are tightly regulated at multiple levels. Recently we have shown that the cyclin C-Cdk8 kinase negatively regulates *ATG8* expression within the Ume6-Rpd3 HDAC axis (Willis *et al*, 2020). This kinase, together with Med13 and Med12 form the Cdk8 kinase module (CKM) of the mediator complex that in yeast predominantly repress transcription of a diverse set of meiosis and stress response genes (Cooper *et al*, 1997; Cooper *et al*, 1999) by interacting with DNA bound transcription factors and RNA polymerase II (Akoulitchev *et al*, 2000; Jeronimo *et al*, 2016; Nemet *et al*, 2014).

Activation of genes controlled by the CKM is achieved by disrupting its association with the mediator (Jeronimo & Robert, 2017). Studies from our group revealed that this is achieved by CKM disassembly. However, we observed that the mechanisms used to disassemble the CKM is dependent upon environmental cues (outlined in Fig. 1) (Cooper *et al*, 2014). In short, oxidative stress triggers cyclin C translocation to the cytoplasm (Cooper *et al*, 2012) where it mediates stress-induced mitochondrial fission and regulated cell death (RCD) in both yeast (Cooper *et al*., 2014) and mammalian cells (Ganesan *et al*, 2019; Jezek *et al*, 2019; Wang *et al*, 2015). Its nuclear release is dependent upon Med13’s destruction by the UPS (Khakhina *et al*, 2014; Stieg *et al*, 2018). In contrast, following a survival cue (nitrogen starvation), cyclin C is rapidly destroyed by the UPS before its nuclear release which prevents mitochondrial fission (Willis *et al*., 2020).

**Figure 1.**
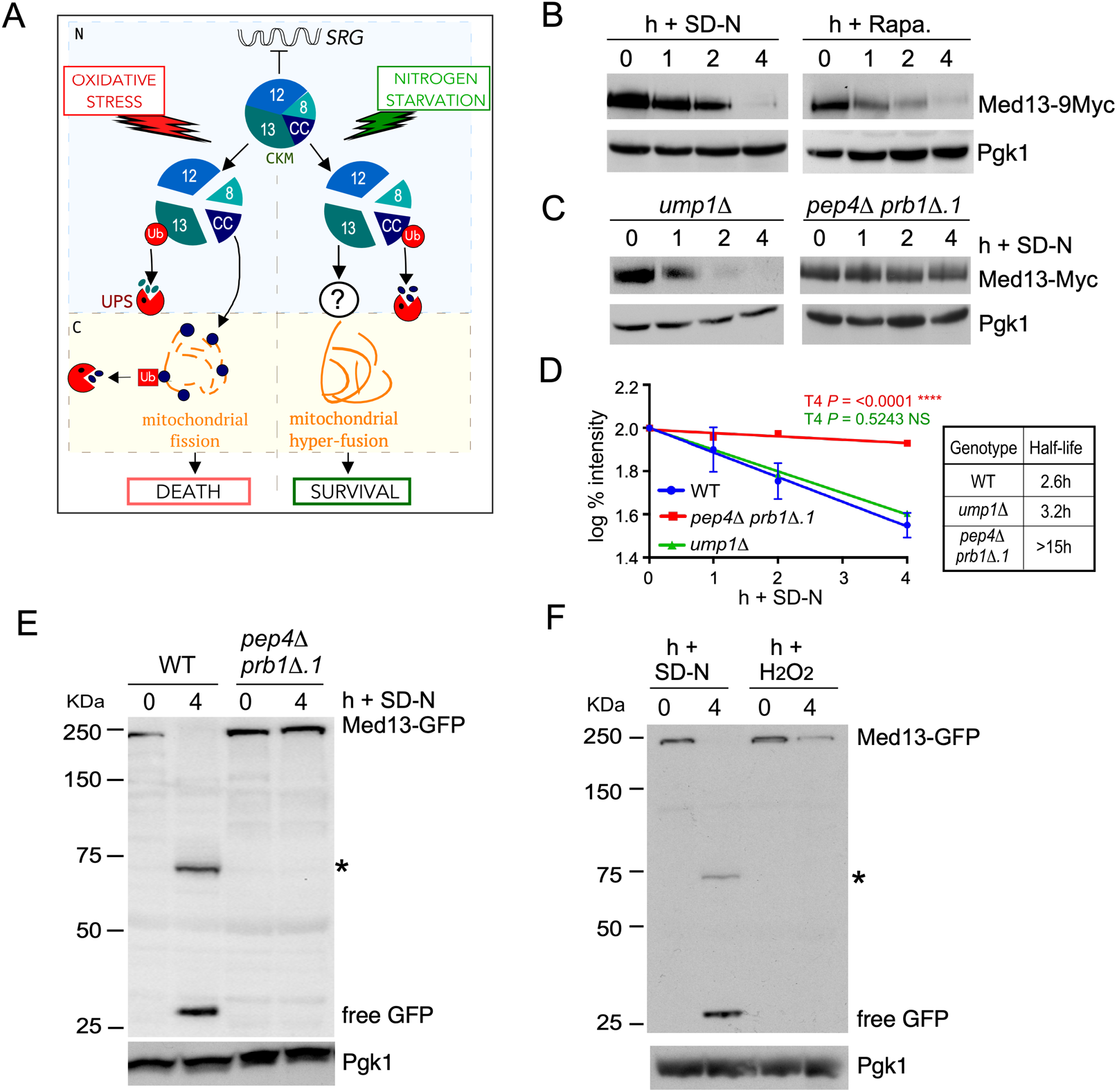
Med13 is degraded via the vacuolar proteolysis following nitrogen starvation. **A** Model outlying how the cyclin dependent kinase module (CKM) of the mediator complex is disassembled following stresses that mediate cell death or survival pathways. Before stress cyclin C (CC), Cdk8 (8), Med13 (13) and Med12 (12) form the CKM that predominantly represses stress response genes (SRGs) (Cooper *et al*., 1997; Cooper *et al*., 1999). Repression is relieved after oxidative stress and nitrogen depletion by CKM disassembly, mediated by different mechanisms. After oxidative stress, Med13 is destroyed by the UPS (Khakhina *et al*., 2014; Stieg *et al*., 2018), which allows cyclin C localization to the mitochondria where it triggers stress-induced mitochondrial fission and promotes cell death (Cooper *et al*., 2014). Following nitrogen starvation, cyclin C is rapidly destroyed by the UPS before its nuclear release to prevent mitochondrial fission (Willis *et al*., 2020) whereas the fate of Med13 is the subject of this current manuscript. N-nucleus, C-cytoplasm. **B** Western blot analysis of extracts prepared from wild-type cells expressing endogenous Med13-9xMyc (RSY2211) resuspended in nitrogen starvation medium (SD-N) or treated with 200 ng/ml rapamycin for the indicated times. **C** As in B except that endogenous Med13 protein levels (13Myc) were monitored in *ump1Δ* (RSY1961) and *pep4Δ prb1Δ.1* (RSY2215) strains. **D** Degradation kinetics and half-life of Med13 protein levels obtained in B and C. Error bars indicate S.D., N=3 of biologically independent experiments. **E** Wild-type (RSY10) or *pep4Δ prb1Δ.1* (RSY449) cells expressing Med13-GFP (pSW218) were starved for nitrogen for indicated times. For all cleavage assays, free GFP refers to the protease resistant GFP moiety that accumulates after the full-length fusion protein is degraded via the vacuole. GFP accumulation was monitored by Western blot analysis using anti-GFP antibodies. An asterisk indicates a nonspecific proteolytic fragment unrelated to autophagy. **F** As in E except that wild-type cells were resuspended in SD-N or treated with 0.8 mM H_2_O_2_. Pgk1 protein levels were used as a loading control.

This study reveals that Med13 is degraded by vacuolar proteolysis by a previously undescribed autophagic pathway which requires the cytosolic nucleoporin Gle1 and the sorting nexin heterodimer Snx4-Atg20. Moreover, two transcriptional activators that regulate *ATG* expression were also degraded upon nitrogen starvation by this mechanism. Taken together, this suggests a model in which Snx4-mediated nucleophagy of *ATG* transcriptional regulators allows fine-tuning of the autophagic response. This highly selective autophagy mechanism rapidly degrades substrates and requires the nuclear pore complex which makes this pathway distinct from previously described nucleophagy pathways.

## RESULTS

### Med13 is actively degraded following nitrogen starvation

We started this investigation by addressing if Med13 was destroyed following nitrogen starvation. Wild-type cells expressing endogenous Med13-9xmyc were starved for nitrogen (SD-N), and Western blot analysis showed that Med13 protein levels decreased with a half-life of 2.6 h (Fig 1B, D, source data Fig 1). Similarly, Med13 was rapidly degraded in replete medium containing rapamycin, a drug that mimics nitrogen starvation by inhibiting TORC1 (Li *et al*, 2014) (Fig 1B, Fig EV1A). As Med13 half-life is >6 h in unstressed cultures (Khakhina *et al*., 2014) and *MED13* mRNA increased following 4 h in SD-N (Fig EV1B), these results indicate that Med13 is actively degraded following TORC1 inhibition.

### Med13 degradation following nitrogen starvation is mediated by the vacuole

Med13 levels were next monitored in *ump1Δ*, a mutant deficient for 20S proteasome assembly (Ramos *et al*, 1998), and no change in degradation kinetics was observed (Fig 1C, D). In contrast, in a vacuolar protease mutant (*pep4Δ prb1Δ.1*) (Takeshige *et al*, 1992; Van Den Hazel *et al*, 1996) Med13 was stable in SD-N (Fig 1C, D) (half-life >15 h) indicating that Med13 degradation requires vacuolar proteolysis. Confirming this, the same results were obtained in wild type, *ump1Δ* and *pep4Δ prb1Δ.1* cells harboring a low copy, functional Med13-3xHA plasmid (Stieg *et al*., 2018) (Fig EV1C, D). Moreover, we found that after nitrogen starvation GFP accumulated in Med13-GFP cleavage assays. This indicates that Med13-GFP is degraded in the vacuole as the compact fold of GFP renders it resistant to vacuolar hydrolases (Shintani & Klionsky, 2004). Accordingly, repeating these cleavage assays in *pep4Δ prb1Δ.1* cells abolished the formation of free GFP and stabilized full-length Med13-GFP (Fig 1E). As anticipated from our previous studies (Khakhina *et al*., 2014) Med13-GFP was destroyed following 0.8 mM H_2_O_2_ and no GFP accumulation was seen (Fig 1F). These results show that the proteolysis machinery employed to degrade Med13 is dependent upon environmental cues.

To visualize Med13 vacuolar degradation we used live-cell imaging of endogenous Med13-mNeongreen in *pep4Δ prb1Δ.1* cells. In SD media, Med13-mNeongreen is nuclear (Fig 2A) but after 4 h in SD-N, Med13 accumulated in the vacuole (Fig 2B). The deconvolved collapsed images in Fig 2C also captured Med13-mNeongreen transitioning between these organelles. After 24 h in SD-N, Med13-mNeongreen is exclusively vacuolar (Fig EV2A), and similar results were obtained when endogenous Med13-YFP was expressed in wild-type cells treated with PMSF that blocks the activity of vacuolar serine proteases (Fig 2B) (Takeshige *et al*., 1992). These results are consistent with a model in which nitrogen starvation triggers Med13 vacuolar proteolysis.

**Figure 2.**
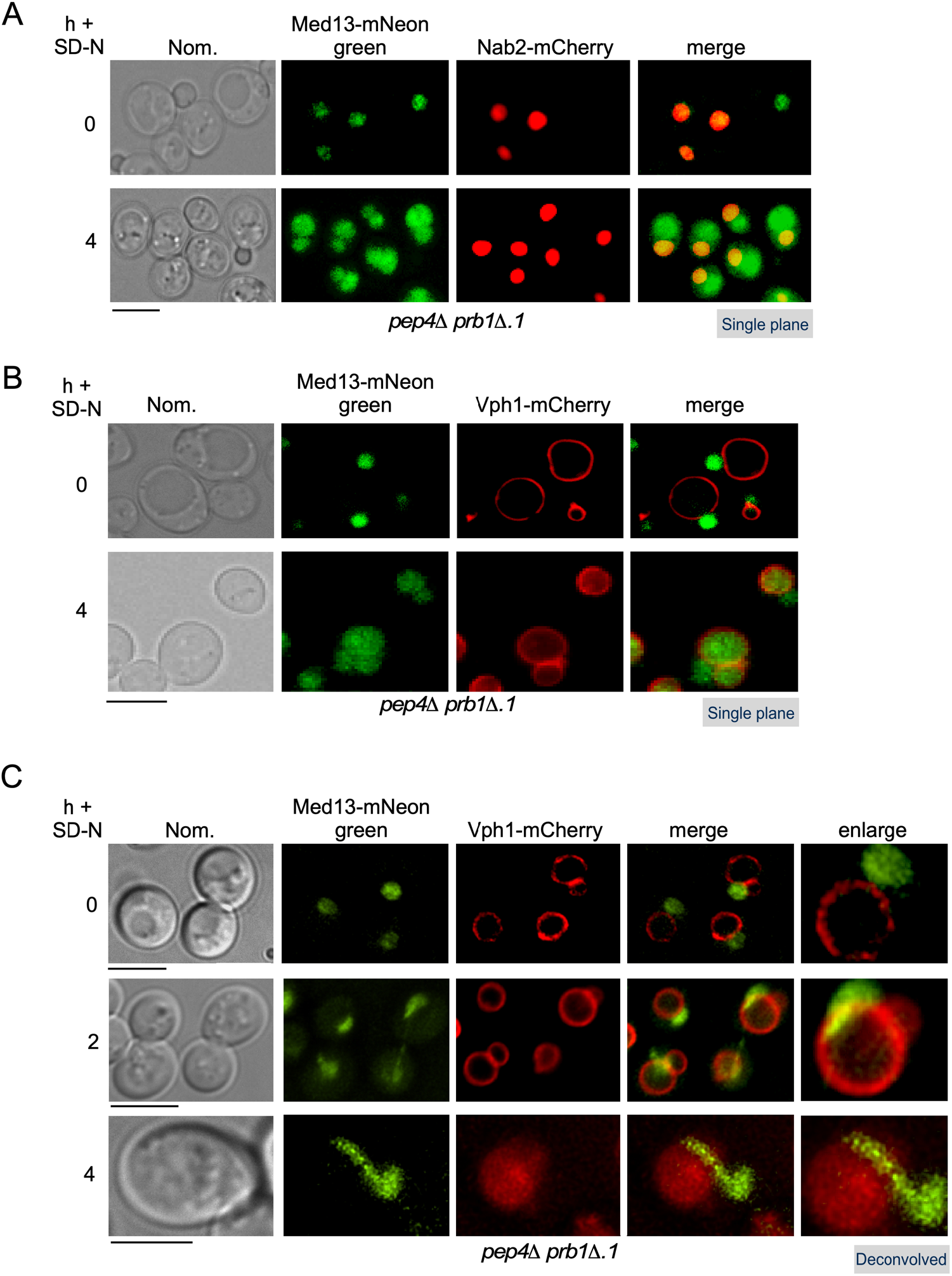
Med13 translocates from the nucleus to the vacuole in nitrogen starvation. **A** Endogenous Med13-mNeongreen localization was monitored in *pep4Δ prb1Δ.1* cells (RSY2305) expressing Nab2-mCherry (a nuclear marker) before (growing in SD) and after 4 h in SD-N. Representative single plane images are shown. **B** As in A, except that cells expressed a vacuolar marker (Vph1-mCherry). **C** As in B, except that the slices were taken through the whole cell which were then collapsed and deconvolved. Representative deconvolved images are shown. Scale bar = 5 µm.

### Med13 degradation requires core autophagy machinery

Next, we investigated if Med13 degradation was dependent upon proteins required for autophagosome biogenesis (see Fig 3A). Med13 was significantly stabilized following nitrogen starvation in mutants defective in induction (*atg1Δ*) or phagophore fusion (*atg8Δ*) (Kamada *et al*, 2000) (half-lives >15 h, Fig 3B, C). Likewise, GFP accumulation from Med13-GFP was abolished in various autophagy mutants (Fig 3A, D). We did observe some turnover of full-length Med13-GFP which we attribute to a combination of Med13 being expressed from the *ADH1* promotor rather than the endogenous locus used for degradation assays, and UPS activity. Consistent with this, Med13-GFP was more stable in *atg1Δ ump1Δ* than *atg1Δ* cells (Fig EV3A). Therefore, both the vacuole and core autophagy proteins are needed for Med13 degradation.

**Figure 3.**
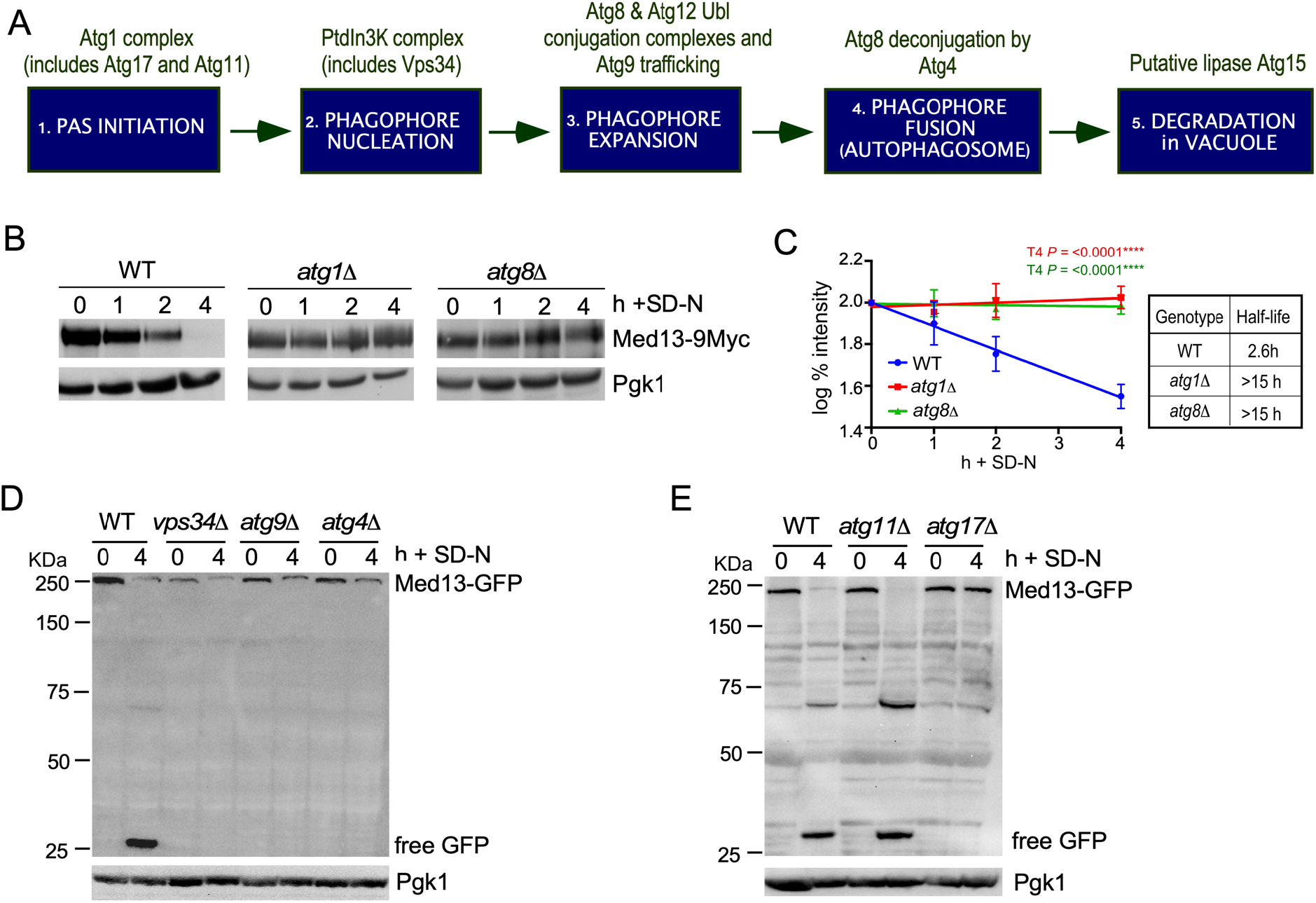
Med13 requires the core autophagy machinery for vacuolar degradation. **A** Schematic depicting the five stages of autophagy and the major Atg protein complexes associated with each stage. **B** Western blot analysis of extracts prepared from wild-type (RSY2211), *atg1Δ* (RSY2214) and *atg8Δ* (RSY2231) cells expressing endogenous Med13-9Myc resuspended in SD-N media for the indicated times. **C** Degradation kinetics and half-life of Med13 protein levels obtained in B. Error bars indicate S.D., N=3 of biologically independent experiments. **D and E** The indicated mutants expressing Med13-GFP (pSW218) were starved for nitrogen for the indicated times and accumulation of free GFP monitored by Western blot analysis using anti-GFP antibodies. An asterisk indicates a nonspecific proteolytic fragment unrelated to autophagy. For all experiments, Pgk1 protein levels were used as a loading control.

### Med13 degradation does not use known selective autophagy pathways

Selective autophagy of excess or damaged cellular components in physiological conditions requires the scaffold protein Atg11 (Zientara-Rytter & Subramani, 2020). In starvation conditions, Atg17 replaces Atg11 and functions as the scaffold protein tethering the Atg1 kinase complex to the growing phagophore (Matscheko *et al*, 2019). Analyzing the autophagic degradation of Med13-GFP (Fig 3E) or degradation of endogenous Med13-9xmyc (Fig EV3B, C) revealed that Atg17 but not Atg11 is required for Med13 destruction. Likewise, known receptors for selective autophagy pathways (Cue5, Atg19, Atg36 and Atg32, Fig EV3D) (Farre & Subramani, 2016) including those needed for micro- and macro-nucleophagy (Nvj1, Atg39 and Atg40, Fig 4A, Fig EV3B, C) (Millen *et al*, 2009; Mochida *et al*., 2015) are also not required. This is consistent with a model that Med13 is degraded via an undescribed nucleophagy pathway.

**Figure 4.**
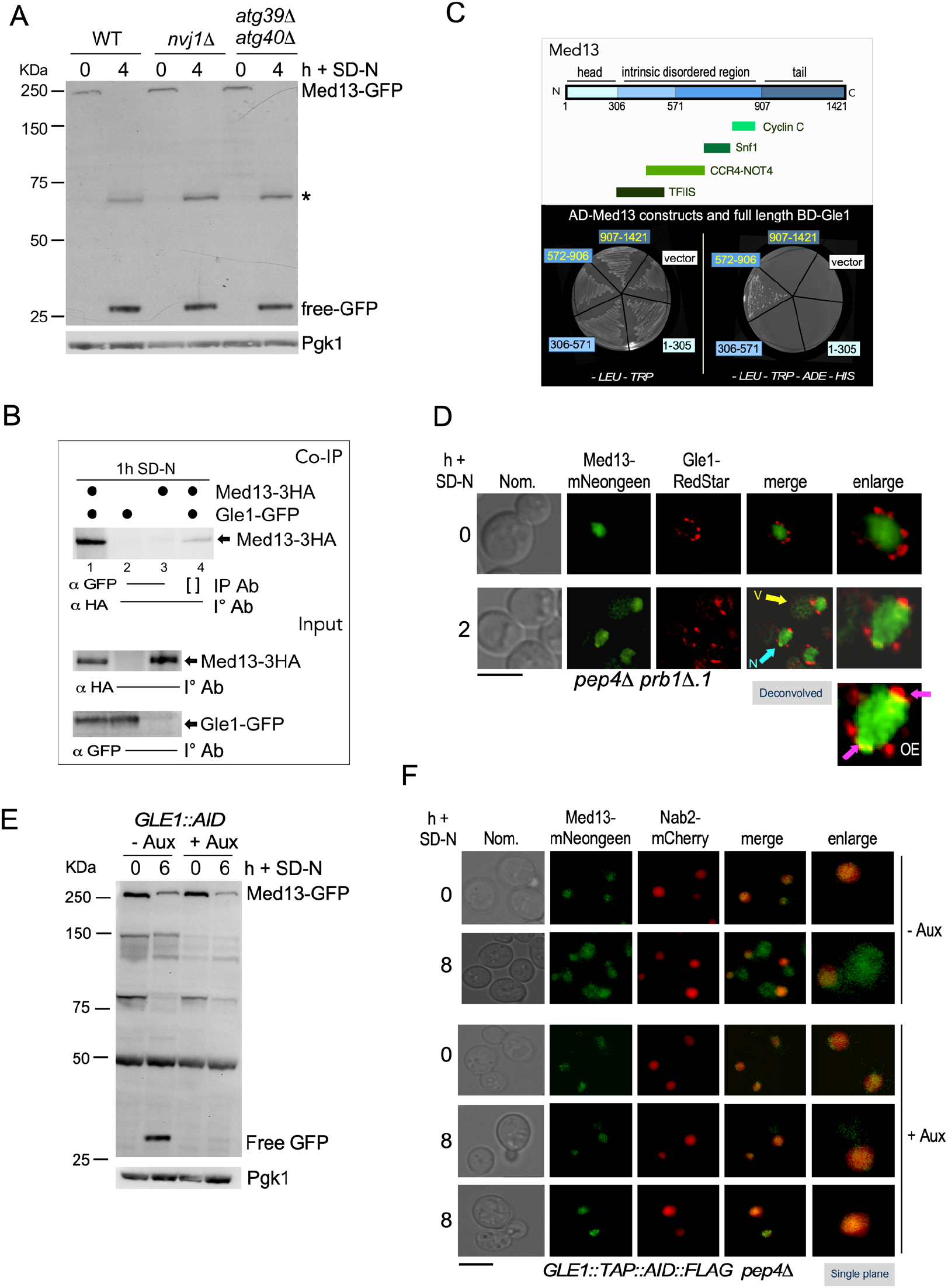
Autophagic degradation of Med13 requires the nucleoporin Gle1 and is independent of known nucleophagy pathways. **A** Western blot analysis of Med13-GFP cleavage assays after 4 h nitrogen depletion in micro-nucleophagy (*nvj1Δ,* RSY2106) and macro-nucleophagy (*atg39Δ atg40Δ,* RSY2123) mutants. An asterisk indicates a nonspecific proteolytic fragment unrelated to autophagy and Pgk1 protein levels were used as a loading control. **B** Co-immunoprecipitation analysis of endogenous Gle1-GFP and Med13-3HA. Whole cell lysates were immunoprecipitated with the antibodies shown from nitrogen-starved *pep4Δ prb1Δ.1* cells expressing endogenous Gle1-GFP (RSY2423) and Med13-3HA (pKC801, lanes 1 and 4) or a vector control (lane 2). *pep4Δ prb1Δ.1* cells expressing Med13-3HA alone (lane 3) was included as a control. [] represents no antibody control. Med13-HA was detected by Western blot analysis of immunoprecipitates. Western blot analysis of the proteins in the whole cell lysates for the three conditions tested is shown (input - bottom panel). **C** Map of Med13 depicting different structural regions and known interacting proteins (upper panel). Different colors represent different regions of the protein and structural regions are denoted by amino acid positions. Med13-Gle1 Y2H analysis. Y2H Gold cells harboring Gal4-BD-Gle1 and the indicated Gal4-AD-Med13 subclone or empty vector control were streaked on medium selecting for plasmid maintenance (left) or induction of the *ADE2* and *HIS3* reporter genes (right) by Y2H interaction (lower panel). See Fig EV5A, B for Western blot analysis of the different constructs. **D** *pep4Δ prb1Δ.1* cells expressing endogenous Med13-mNeongreen and endogenous Gle1-RedStar (RSY2450) were starved for nitrogen for 2 h and monitored by fluorescence microscopy. Deconvolved representative images are shown. The yellow arrow is pointing to Med13-GFP in the vacuole whereas the blue arrow shows co-localization. OE represents an over-exposed image to better show the colocalization of Gle1and Med13 (pink arrows) Scale = 5μm. **E** Med13-GFP cleavage assays performed in the Gle1 Auxin-inducible degron (Gle1-AID) strain (RSY2456). Cells expressing Med13-GFP (pSW320) were treated with 250 μM Auxin for 30 m before proceeding with autophagic cleavage assays in SD-N. Free GFP accumulation was then detected using Western blot analysis. **F** Fluorescence microscopy of Med13-mNeongreen localization in the Gle1 Auxin-inducible degron (Gle1-AID) strain expressing Nab2-mCherry before and after SD-N. An 8 h time point was used as the strain background is BY4741 which is not as sensitive to environmental stress as W303a. Scale = 5 µm.

### The cytoplasmic nucleoporin Gle1 associates with Med13 after nitrogen starvation

To define components of this pathway, pull-down assays using Med13-3xHA followed by mass-spectroscopy was used (Fig EV4A, B and Table I for a list of candidate interactors). The conserved, essential nucleoporin Gle1, that localizes to the cytoplasmic filament of the nuclear pore complex (NPC) was identified. Gle1 is a regulator of DEAD-box proteins that are required to facilitate changes to ribonucleoprotein complexes involved in mRNA export and translation initiation (Alcazar-Roman *et al*, 2006; Aryanpur *et al*, 2017; Weirich *et al*, 2006). Co-immunoprecipitation analysis revealed that Med13-3xHA and endogenous Gle1-GFP interacted following 1 h SD-N, confirming this interaction (Fig 4B, source data Fig 2). Yeast two-hybrid assays (Y2H) showed that the large central intrinsic disordered region (IDR) of Med13, which provides a flexible interaction hub for multiple partners (Stieg *et al*., 2018; Uversky, 2011), interacts with Gle1 (Fig 4C, Fig EV5A, B). In addition, endogenous Med13-mNeongreen and Gle1-RedStar were followed by live-cell imaging. Med13 moves from being diffuse nuclear to associating with the punctate Gle1-RedStar that surrounds the nucleus on its passage to the vacuole after nitrogen starvation (Fig 4D).

Gle1 is an essential protein (Murphy & Wente, 1996), therefore, the auxin-inducible degron (AID) system was used to reduce Gle1 protein levels (Fig EV4C). The autophagic degradation of Med13-GFP was significantly decreased following auxin treatment (Fig 4E) as monitored by GFP accumulation. Moreover, after nitrogen depletion, Med13-mNeongreen was retained in the nucleus in the *GLE1*-degron (Fig 4F). However, Crm1 and Msn5, two major Ran-dependent β karyopherins that export mRNA and nuclear proteins following stress (Hutten & Kehlenbach, 2007; Mosammaparast & Pemberton, 2004; Yoshida & Blobel, 2001), are not required for Med13 autophagic degradation (Fig EV4D, E). This strongly suggests that Med13 is transported through the NPC to exit the nucleus, ending at the Gle1 exchange platform.

### The sorting nexin-heterodimer Snx4-Atg20 is required for efficient autophagic degradation of Med13

To further define components of this new nucleophagy mechanism, null alleles of candidate proteins from the mass spectroscopy were screened using Med13-GFP cleavage assays. In cells deleted for the conserved sorting nexin Snx4 (Atg24), we found that free GFP was significantly reduced (Fig 5A). Snx4 is a member of the conserved sorting nexin family of proteins. Snx4 forms distinct heterodimer complexes, with either Snx41 or Atg20, which mediate retrograde trafficking of cargo from the vacuole and endosomes to the Golgi. This role maintains homeostasis and is dispensable for non-selective autophagy (Ma & Burd, 2020; Ma *et al*, 2017; Ma *et al*, 2018; Suzuki & Emr, 2018). In contrast, Snx4-Atg20 is essential for many selective autophagy pathways including mitophagy and pexophagy (Deng *et al*, 2013; Kanki *et al*, 2009; Nice *et al*, 2002; Popelka *et al*, 2017; Shpilka *et al*, 2015)

**Figure 5.**
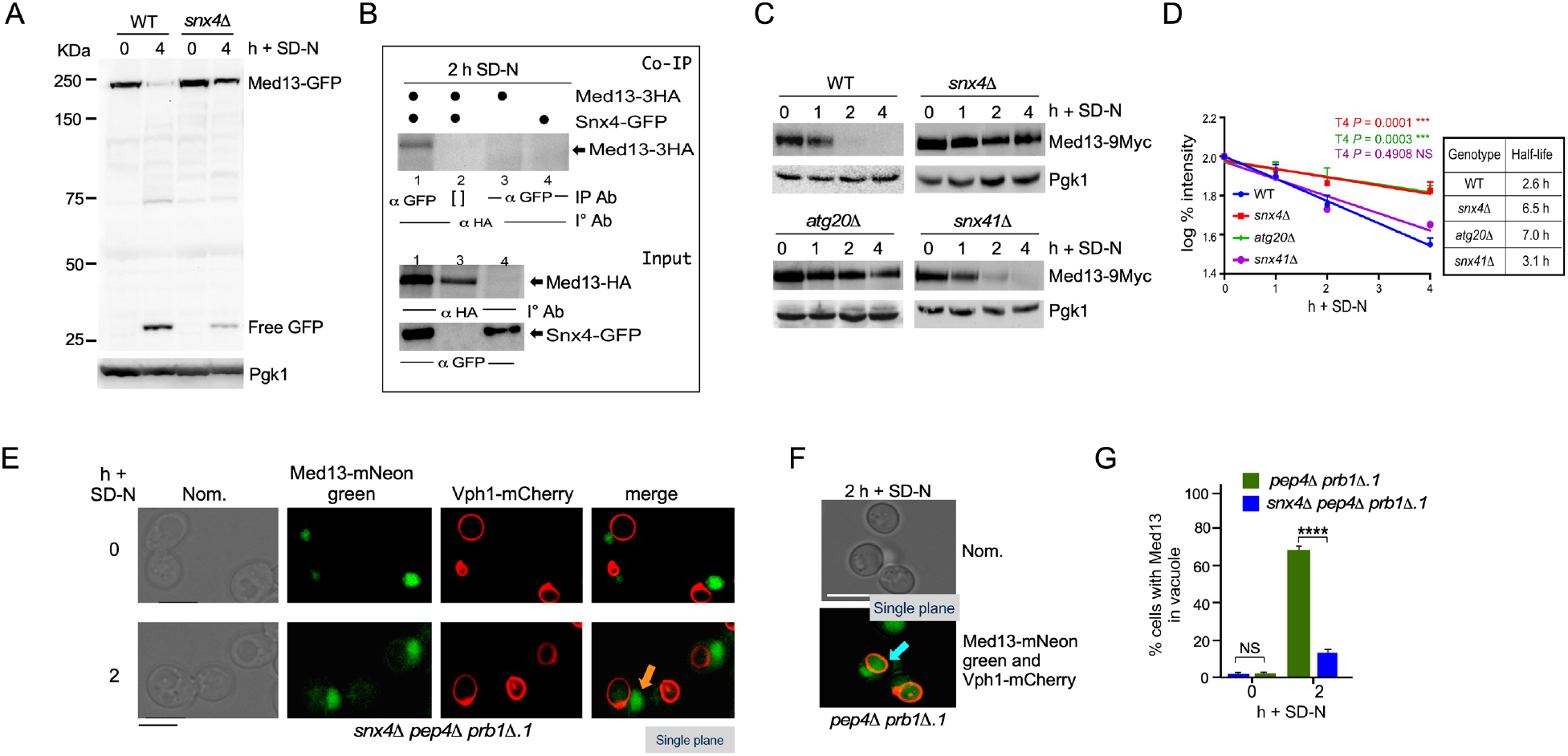
The sorting nexin heterodimer, Snx4-Atg20 is required for efficient autophagic degradation of Med13. **A** Western blot analysis of Med13-GFP cleavage assays after 4 h nitrogen starvation in wild-type and *snx4Δ* (RSY2272). **B** Co-immunoprecipitation analysis of GFP-Snx4 and Med13-3HA. Whole cell lysates were immunoprecipitated with the antibodies shown from nitrogen-starved *pep4Δ prb1Δ.1* cells expressing GFP-Snx4 (RSY2299) and Med13-3HA (pKC801, lanes 1 and 2) or a vector control (lane 4). *Pep4Δ prb1Δ.1* cells expressing Med13-3HA alone (lane 3) was included as a control. [] represents no antibody control. Med13-HA was detected by Western blot analysis of immunoprecipitates. Western blot analysis of the proteins in the whole cell lysates for the three conditions tested is shown (input - bottom panel). **C** Western blot analysis of extracts prepared from wild-type (RSY2211), *snx4Δ* (RSY2276), *atg20Δ* (RSY2277) and *snx41Δ* (RSY2394) expressing endogenous Med13-9Myc resuspended in SD-N for the indicated times. Pgk1 was used as a loading control. **D** Degradation kinetics and half-lives of Med13 protein levels obtained in B. Error bars indicate S.D., N=3 of biologically independent experiments. **E** Fluorescence microscopy of endogenous Med13-mNeongreen localization in *snx4Δ pep4Δ prb1Δ.1* (RSY2324) expressing the vacuole marker Vph1-mCherry. Cells were visualized before (SD) and after 2 h of SD-N treatment and representative single plane images are shown. Scale = 5 µm. **F** As in D except that endogenous Med13-mNeongreen localization was followed in nitrogen-starved *pep4Δ prb1Δ.1* cells. Representative single plane images of the results are shown. Bar = 5µm. **G** Quantification of accumulation of Med13-mNeongreen in vacuoles obtained from results in D and E. 100 cells counted per sample. N=3 biological samples. *\*\*\*\* P* = >0.0001.

Co-immunoprecipitation analysis confirmed the interaction between Med13 and Snx4 in SD-N (Fig 5B). Med13 degradation assays revealed that Snx4 and Atg20, but not Snx41, mediate Med13 autophagic degradation (Fig 5C, D). The half-life of Med13 in these mutants was 6.5 and 7.0 h, respectively, compared to >15 h seen in core autophagic mutants. Cytosolic and vacuolar Med13-mNeongreen levels were drastically reduced in *snx4Δ pep4Δ prb1Δ.1* mutants compared to the *pep4Δ prb1Δ.1* control (Fig 5E, F, G). These data demonstrate that the Snx4-Atg20 heterodimer is required for maximal Med13 autophagic degradation, but in its absence, limited autophagic degradation of Med13 still can occur. Taken together these results show that Snx4-Atg20 heterodimer mediates the efficient autophagic degradation of Med13

### Snx4 localizes to the nuclear periphery to transport Med13 to autophagosomes

Snx4 binds to the scaffold protein Atg17 (Nice *et al*., 2002), whose major autophagic role is binding the Atg1 kinase to the PAS (Hollenstein & Kraft, 2020). As both Snx4 and Atg17 mediate Med13 degradation, this suggests a model in which once Med13 passes through the NPC, it is recognized by Snx4 and delivered to the growing phagophore by Snx4-Atg17 association. Consistent with this, Med13-3xHA and endogenous Atg17-GFP co-immunoprecipitated following 2 h in SD-N. In *snx4Δ,* this interaction was drastically decreased (Fig 6A). Also, endogenous Atg17-RedStar co-localized with Med13-mNeongreen after nitrogen starvation (Fig EV6A). This suggests that Snx4-Atg17 interaction promotes the efficient recruitment of Med13 to autophagosomes. Quantitative co-localization analysis with GFP-Snx4 with the nuclear marker Nab2-mCherry, showed that nitrogen depletion triggers a ∼10-fold increase in perinuclear GFP-Snx4 foci (Fig 6B, Fig EV6B, C). Moreover, Med13-mNeongreen co-localized with both perinuclear and cytosolic Snx4 foci in SD-N (Fig 6C and EV6D). Together, this suggests that Snx4 localizes to the nuclear periphery to retrieve and transport Med13 to autophagosomes via Atg17 association.

**Figure 6.**
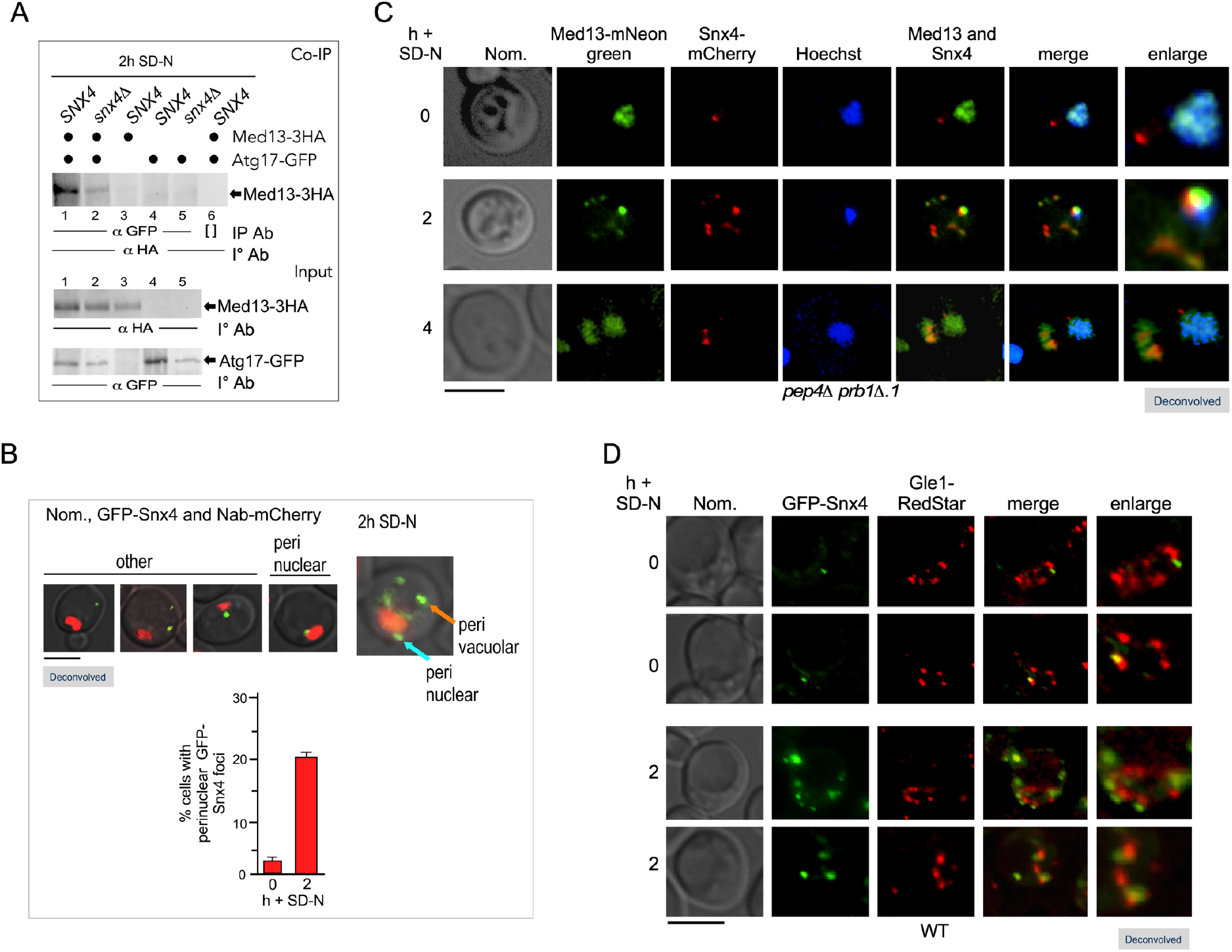
Snx4 localizes to the nuclear periphery to retrieve Med13. **A** Co-immunoprecipitation analysis of endogenous Atg17-GFP and Med13-3HA in the presence and absence of Snx4. Whole cell lysates were immunoprecipitated with the antibodies shown from nitrogen-starved *pep4Δ prb1Δ.1* (RSY2395) *or snx4Δ pep4Δ prb1Δ.1* cells (RSY2396) expressing endogenous Atg17-GFP and Med13-3HA (pKC801, lanes 1, 2 and 6) or a vector control (lanes 4 & 5). *Pep4Δ prb1Δ.1* cells expressing Med13-3HA alone (lane 3) was included as a control. [] represents no antibody control. Med13-HA was detected by Western blot analysis of immunoprecipitates. Western blot analysis of the proteins present in the whole cell lysates for the conditions tested is shown (input - bottom panel). **B** Representative images showing perinuclear and perivacuolar localization of GFP-Snx4 in wild-type expressing Nab2-mCherry (nuclear marker) before and after nitrogen starvation. Bar = 5μm. The number of perinuclear foci was counted (N=2) before and after nitrogen starvation. At least 100 cells were counted per sample. Data are the percentage of perinuclear foci among the total number of foci. Scale = 5 µm. **C** Fluorescence microscopy of *pep4Δ prb1Δ.1* cells expressing endogenous Med13-mNeongreen and mCherry-Snx4 (RSY2424) before and after nitrogen depletion. Hoechst staining was used to visualize the nucleus. Representative deconvolved images are shown. Bar = 5μm. **D** GFP-Snx4 and endogenous Gle1-RedStar co-localize in wild-type cells (RSY2451) following nitrogen starvation. Representative images are shown. Bar = 5μm.

The above model predicts that Snx4 and Gle1 interact whilst “handing-off” Med13 from the NPC to the sorting nexin complex. Live-cell imaging showed that perinuclear Snx4 foci are adjacent to, or co-localize with, Gle1 in unstressed cultures. After nitrogen starvation, an increased number of perinuclear Snx4 foci co-localize with Gle1 (Fig 6D and Fig EV6E), which is consistent with our model that Snx4 localizes to the NPC to retrieve Med13.

### Snx4 specifically targets for Med13 autophagic degradation

To further understand the sequential stages of this selective autophagy pathway, we next asked if Med13 nuclear localization was required for retrieval and transport by Snx4. To address this, we fused Med13-GFP to the N-terminus of Crn1, a protein associated with actin rafts, which relocalized Med13 to the plasma membrane (Humphries *et al*, 2002) (Fig 7A). Autophagic degradation assays with Crn1-Med13-GFP showed that GFP accumulated in wild-type cells, and this was mostly dependent on Snx4 (Fig 7B). This confirms that Med13 can be targeted for Snx4-mediated autophagic destruction even when located outside the nucleus. Autophagic degradation of Pgk1-GFP, an established substrate of non-selective autophagy (Welter *et al*, 2010), is Snx4 independent (Fig EV7A). This provides further evidence that Snx4 binds to Med13 after nuclear release and this interaction is highly specific.

**Figure 7.**
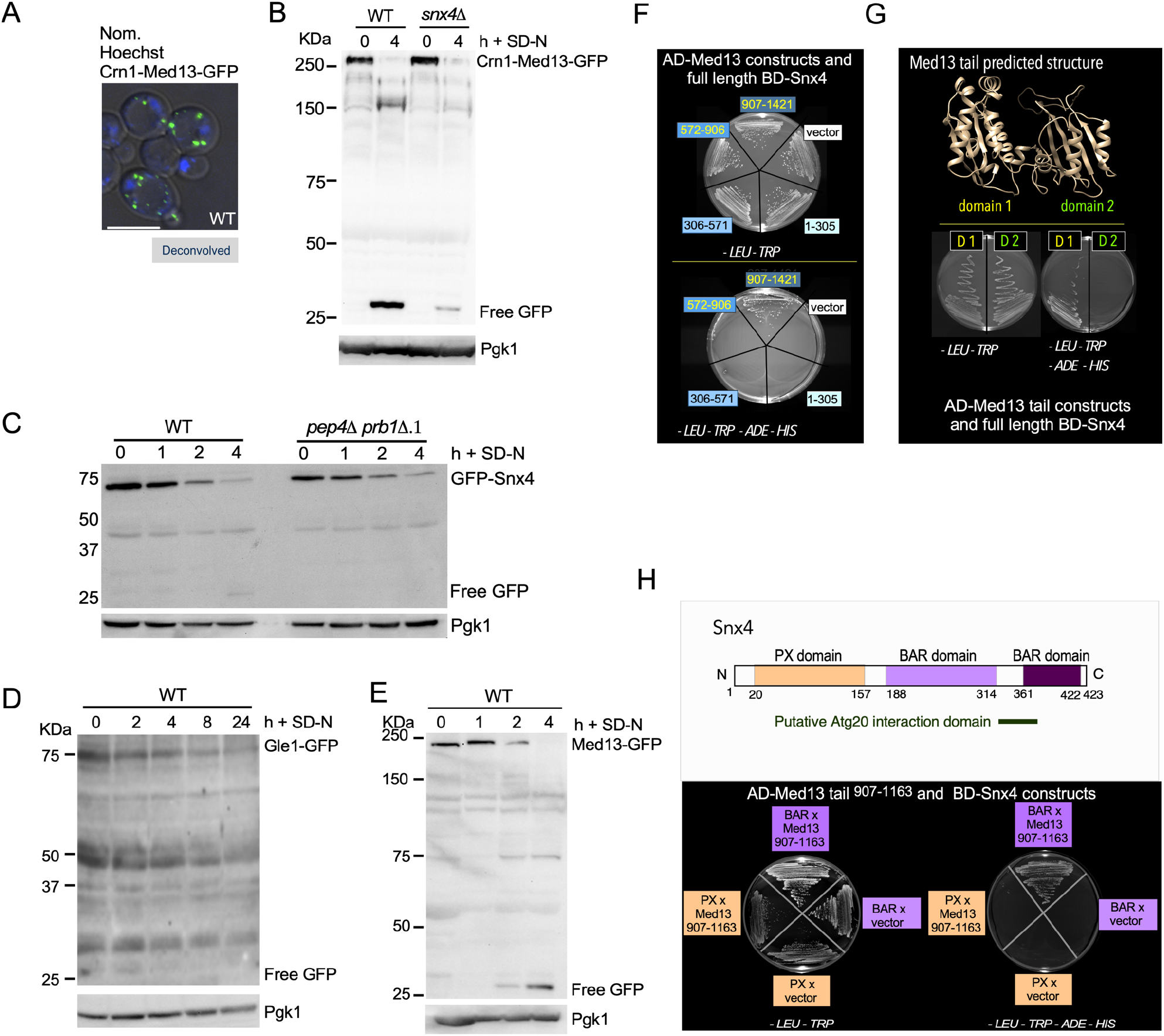
The BAR domain of Snx4 interacts with the C-terminal region of Med13. **A** Fluorescence microscopy of Crn1-Med13-GFP (pSW288) in wild-type cells growing in SD. Hoechst staining was used to visualize the nucleus. Bar = 5μm. **B** Western blot analysis of Med13-GFP (pSW218) or Crn1-Med13-GFP (pSW288) cleavage assays performed in wild-type or *snx4Δ* cells following nitrogen starvation. **C** Western blot analysis of GFP-Snx4 cleavage assays performed in wild-type (RSY2283) and *pep4Δ* (RSY2299) cells following nitrogen starvation. **D** Western blot analysis of Gle1-GFP cleavage assays performed in wild-type (RSY2455) cells following prolonged nitrogen starvation (24 h). **E** Western blot analysis of Med13-GFP (pSW218) cleavage assays performed in wild-type cells. Pgk1 was used as a protein loading control for all experiments. **F** Y2H Gold cells harboring Gal4-BD-Snx4 and the indicated Gal4-AD-Med13 construct or vector control were plated on medium selecting for plasmid maintenance *(-LEU, -TRP*) (top) or interaction by induction of the *ADE2* and *HIS3* reporter genes (bottom). See Fig EV5A, B for Western blot analysis of the different constructs. **G** Predicted structural analysis of the Med13 tail region using Phyre2 plot analysis of this region (Kelley *et al*., 2015). Y2H analysis of full-length Gal4-BD-Snx4 and Gal4-AD-Med13 subclones containing either the first or second domain region of the Med13 tail. Cells were streaked on medium selecting for plasmid maintenance (left) or induction of reporter genes (right) by Y2H interaction. **H** Map of Snx4 depicting known domains (left panel). Y2H analysis of Snx4 PX and BAR binding domain constructs with the Med13^907-1163^AD construct and empty vector. Cells were streaked on medium selecting for plasmid maintenance (left) or induction of reporter genes (right) by Y2H interaction (right panel).

We next addressed whether Gle1 and Snx4 remain part of the Med13 complex delivered to the vacuole. No substantial accumulation of GFP was seen from GFP-Snx4 (Fig 7C) or Gle1-GFP (Fig 7D), suggesting that neither protein is incorporated into autophagosomes or degraded by the vacuole. Importantly, by monitoring the timing of GFP accumulation from Med13-GFP, we observed that the earliest detection of GFP occurred at 2 h of nitrogen depletion when both Gle1 and Snx4 are present (Fig 7E). These data support the model that Gle1 and Snx4 mediate Med13 delivery to the autophagosome, but these proteins themselves are not vacuolar substrates.

### Snx4 BAR domains interact with the C-terminal domain of Med13

To further understand the interaction between Gle1, Snx4, and Med13, we used Y2H analysis to ask if Snx4 and Gle1 associates with the same or different domains of Med13. The results show that Snx4 interacts with the structured C-terminal tail domain of Med13 (Fig 7F, EV5C). Phyre2 plot analysis of this region (Kelley *et al*, 2015) revealed two potential domains (Fig 7G), of which only one, Med13^907-1163^ interacted with Snx4 (Fig 7G). Therefore Snx4 interacts with a previously undescribed region of Med13 that lies adjacent to the Gle1 interaction domain. Moreover, as Snx4 is not a nuclear protein, this interaction may be direct and define a new role for Snx4 in transporting nuclear proteins.

Snx4 is a conserved member of the SNX-BAR subfamily of sorting nexin proteins. Common to all SNX family members it contains a phosphoinositide-binding phox homology (PX) domain, which binds to phosphatidylinositol 3-phosphate enriched endosomal membranes. It also contains two BAR (Bin/|Amphiphysin/Rvs) domains that bind to curved membranes upon dimerization (Popelka *et al*., 2017; Stanishneva-Konovalova *et al*, 2016). Y2H interaction analyses between either Snx4 domains (PX or BAR domains) and Med13^907-1163^ indicated that the Med13-Snx4 interaction occurs through the BAR domain region (Fig 7H). Taken together, these results indicate that the BAR domains recognize Snx4 cargo binding as well as dimerization partners.

### Med13 negatively regulates the transcription of a subset of *ATG* genes

We next explored if this newly described nucleophagy pathway affects viability during nitrogen starvation. Consistent with previous studies (Nemec *et al*, 2017), efficient survival in prolonged nitrogen starvation conditions requires Snx4 (Fig EV7B). Snx4 is dispensable for non-selective autophagy but required for many forms of non-selective autophagy (Ma *et al*., 2018). This suggests that the role of Snx4 in these pathways is essential for cellular adaptation and survival in prolonged starvation conditions. Survival during periods of nutrient depletion requires the upregulation of *ATG* genes. As Med13 destruction following H_2_O_2_ relieves repression on SRGs (Khakhina *et al*., 2014), here asked if a similar strategy was used for the nitrogen starvation response. RT-qPCR analysis of *ATG* mRNA levels in unstressed wild-type and *med13Δ* cells showed a ∼fourfold increase in *ATG8* mRNA levels (Fig 8A) that was mirrored by Atg8 protein levels (Fig EV7C). Furthermore, *ATG1* and *ATG14* mRNA levels were also elevated (Fig 8A), indicating Med13 represses transcription of a subset of *ATG* genes. Moreover, they are consistent with the model that the destruction of both cyclin C and Med13 following nitrogen starvation are mechanisms used to relieve this repression. These results indicate that Snx4 mediates a new specialized autophagy program able to target a transcriptional repressor that provides a positive feedback loop for the autophagic response.

**Figure 8.**
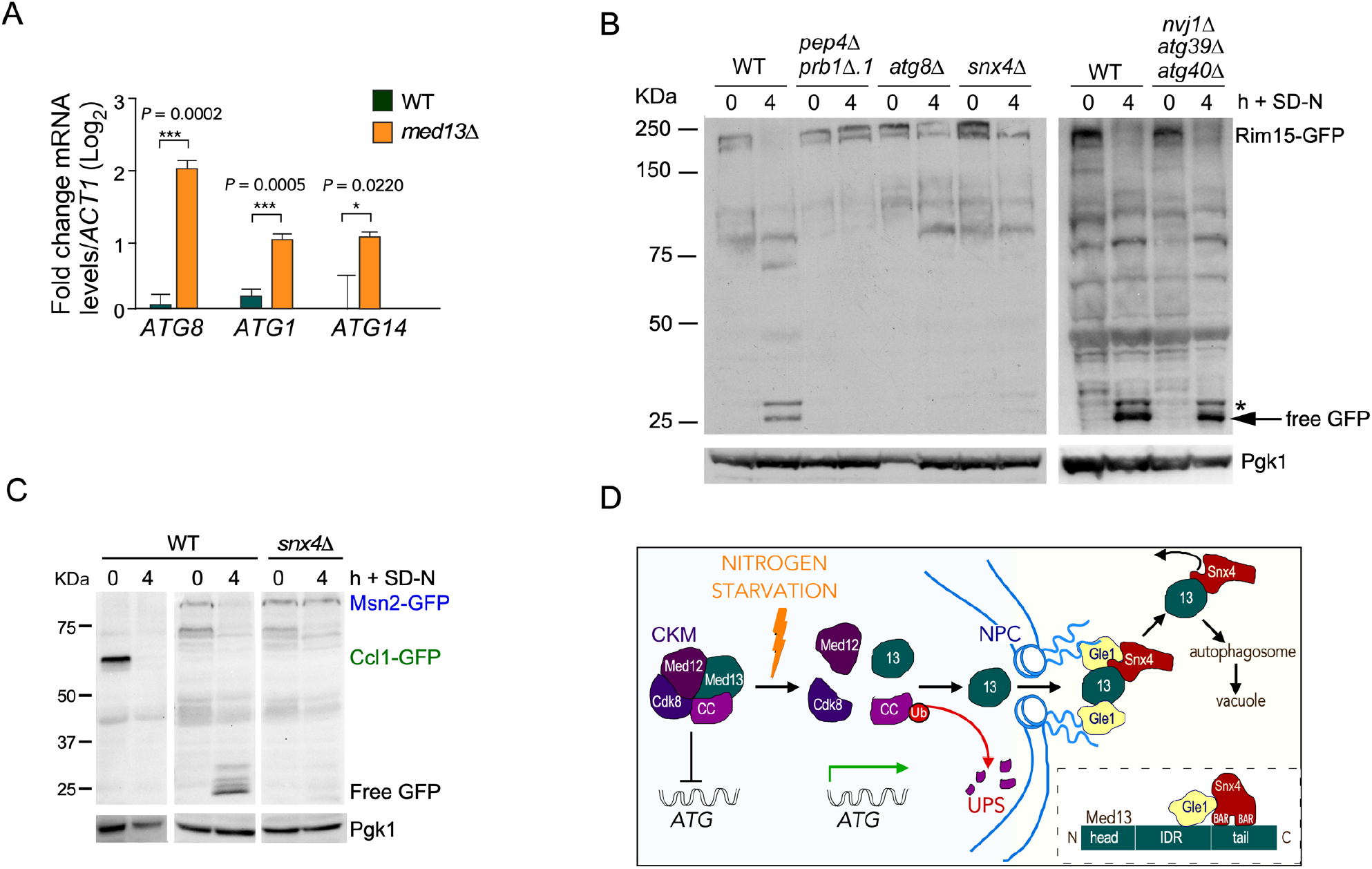
Transcription factors regulating *ATG* genes are nucleophagy substrates. **A** RT-qPCR analysis probing for *ATG8*, *ATG1* and *ATG14* mRNA expression in wild-type and *med13Δ* (RSY2444) cells in unstressed conditions. ΔΔCt results for relative fold change (log_2_) values using wild-type unstressed cells as a control. Transcript levels are given relative to the internal *ACT1* mRNA control. **B** Western blot analysis of Rim15-GFP (pFD846) cleavage assays in indicated mutants after nitrogen starvation. The asterisk denotes a background band. **C** Western blot analysis of Ccl1-GFP (pSW230) and Msn2-GFP (pSW217) cleavage assays in wild-type or *snx4Δ* cells. Pgk1 protein levels were used as a loading control in all experiments. **D** Summary model of novel Snx4-mediated nucleophagy mechanism that selectively targets transcription factors for autophagic degradation. In unstressed cells the CKM represses a subset of *ATG* genes. Following nitrogen starvation an unknown signal triggers the CKM to disassembly. Med13 is transported by unknown mechanisms through the NPC to the cytoplasmic nucleoporin Gle1. Gle1 releases Med13 to the Snx4-Atg20 heterodimer where it interacts with the Snx4 BAR domains (see insert), which in turn interacts with the Atg17 scaffold protein found on growing PAS structures.

### Snx4-mediated nucleophagy degrades other transcription factors following nitrogen starvation

Next, we wanted to explore the idea that Snx4-mediated nucleophagy is a mechanism used by the cell to fine-tune the autophagic response at the level of transcription. We first examined the transcriptional activator Rim15 that also regulates *ATG* genes. During nitrogen starvation, Rim15 enters the nucleus to directly phosphorylate and inhibit the activity of the transcriptional repressors Ume6 (Bartholomew *et al*, 2012) and Rph1 (Bernard *et al*, 2015). Surprisingly, Rim15-GFP cleavage assays revealed Rim15 degradation by Snx4-dependent vacuolar proteolysis that is independent of known nucleophagy pathways (Fig 8B).

To address if Snx4-mediated nucleophagy may have a more global role in degrading transcription factors that control *ATG* expression, the transcriptional activators Msn2 and Ccl1 were tested (Bernard *et al*., 2015; Vlahakis *et al*, 2017; Zhu *et al*, 2016). Free GFP accumulated only from Msn2-GFP in wild-type cells following nitrogen starvation, illustrating that Msn2 is degraded via the vacuole, and this degradation requires Snx4 (Fig 8C). These results support the conclusion that Snx4-mediated degradation of transcription factors targets a unique subset of nuclear proteins. These findings also provide the first evidence that the autophagic pathway directly targets regulatory proteins that control its own processes.

## DISCUSSION

Here, by following Med13’s fate after nitrogen starvation, we have uncovered a previously undescribed nucleophagy pathway in which both negative and positive transcriptional regulators of *ATG* genes are degraded. Our findings, together with previous studies, suggest a two-step pathway involving the NPC and Snx4 for Med13 translocation from the nucleus to the autophagic machinery (outlined in Fig 8D). First, Med13 disassociates from the CKM, shuttles through the NPC and associates with the cytoplasmic nucleoporin Gle1. In the second step, Med13 is handed-off from Gle1 to the sorting nexin heterodimer Snx4-Atg20. Once assembled, this complex localizes to the PAS via the non-selective autophagy scaffold Atg17. Lastly, Snx4-Atg20 is recycled back to the cytosol, and Med13 is degraded by vacuolar proteolysis. This new pathway is distinct from previously identified mechanisms, being the first described pathway to specifically target transcription factors for autophagic degradation. This mode of degradation does not require the canonical nucleophagy mechanisms but instead requires the NPC. Notably, the pathway uses the non-selective scaffold Atg17, and lastly the process is rapid, occurring within 4 h making it distinct from other forms of selective autophagy which require longer starvation times.

How does Med13 find its way to the autophagosome? Med13 is a large, 160 kDa protein that must require an active process to transit from the nucleus to the cytoplasm. Its unusual structure, two folded domains separated by a ∼1000 amino acid IDR region, suggests it functions as a scaffold and interaction hub. This model is supported by both physical (Nagulapalli *et al*, 2016) and genetic studies (Stieg *et al*., 2018). Consistent with this activity, the IDR of Med13 interacts with Gle1, cyclin C, and Cdk8 (see Fig. 4) and is the target of additional regulatory protein kinases including Snf1 (Willis *et al*, 2018). The established role of the Gle1-Dbp5 complex in releasing RNPs (ribonucleoproteins) from mRNPs (messenger ribonucleoprotein particles) (Folkmann *et al*, 2011) sets a precedent for Gle1 handing off proteins once they transit through the NPC. Moreover, recent structural data places the Gle1-Dbp remodeling complex right over the NPC’s central channel, allowing it to efficiently capture proteins as they reach the cytoplasmic side of the NPC (Fernandez-Martinez *et al*, 2016). Although speculative, a similar capture and release mechanism may be used in the Med13 handoff between Gle1 to Snx4. In support of this model, Gle1 and Snx4 co-localize at the NPC (Fig. 6D), but currently, it is unknown if these proteins directly interact.

The best-characterized cargo of Snx4-Atg20 is Snc1, a plasma membrane-directed v-SNARE, but it remains unknown how this protein binds to Snx4 (Bean *et al*, 2017; Hettema *et al*, 2003; Ma *et al*., 2017). Similar to our results with Med13, Snc1 associates with the BAR domain of Snx4 using both co-immunoprecipitation analysis as well as Y2H assays (Zhang *et al*, 2009). Taken together, these results suggest that the BAR domain of Snx4 can both recognize cargos as well as be used for dimerization with Atg20. As the newly defined vital function of autophagy receptor proteins is to stably connect autophagy cargo with scaffold proteins (Hollenstein *et al*, 2019) our results place Snx4 as a receptor protein for this novel nucleophagy mechanism. Supporting this, Snx4 co-localizes with Med13 at both perinuclear and cytoplasmic locations. Our finding that efficient Med13 and Atg17 co-immunoprecipitation requires Snx4 suggests that this sorting nexin relays Med13 from the NPC to the autophagic machinery. In addition, Snx4 mediates Med13 vacuolar degradation when this nuclear protein is relocalized to the plasma membrane, illustrating that Snx4 specifically targets and transports Med13 to growing autophagosomes. Defining the Snx4 interaction motif on Med13, as well as Msn2 and Rim15, may result in a common “Snx4 recognition motif” that could be used to identify additional transcription factors that are degraded by this mechanism. This would be useful because, despite the recent discovery of additional Snx4 cargos, (Bean *et al*., 2017) no consensus Snx4-dependent sorting signal has been identified (Ma & Burd, 2020). This is important as Snx4 is evolutionarily conserved (Zhang *et al*, 2018), with recent discoveries that Snx4 dysregulation is now associated with the etiology of many diseases, including, cancer, Parkinson’s disease and Alzheimer’s disease (Hu *et al*, 2015).

The most unexpected finding from this research was that the complex process of autophagy is used to degrade transcription factors. More unexpectedly, was that two transcriptional activators of *ATG* genes are also regulated by this pathway. This suggests that nucleophagy can fine-tune transcription and points to a new regulatory role for autophagy beyond its canonical function of recycling damaged/unnecessary proteins or organelles. The molecular details of how these transcription factors are initially targeted and then transported through the NPC are unknown. These details will define the nuclear components required for this pathway. This is important as nucleophagy is the least well defined of all the autophagy pathways and in recent years links between deficiencies and human diseases are starting to emerge (Papandreou & Tavernarakis, 2019). Given the highly conserved nature of both Med13 and Snx4 (Tsai *et al*, 2013; Zhang *et al*., 2018), these studies are likely to be relevant to mammalian systems.

## METHODS

### Yeast strains and plasmids

Experiments were primarily performed with endogenously labeled proteins in the *S. cerevisiae* W303 background (Ronne & Rothstein, 1988) and are listed in Table S2. All strains were constructed using replacement methodology (Janke *et al*, 2004). Other strains used were from the Research Genetics yeast knock out collection (Chu & Davis, 2008) and are derived from BY4741 strain background. The Y2H assays were performed in the Y2H Gold strain (PT4084-1, Takara 630489, Matchmaker Gold Yeast Two-Hybrid System). The *GLE1*-Auxin-inducible depletion strain (Fig 4E, RSY2456) was a gift from K. Cunningham (Snyder *et al*, 2019). Live cells were treated with 250 μM Auxin (Indole-3-acetic acid, Gold Bio I-110) dissolved in ethanol 30 m before nitrogen starvation. The doxycycline-inducible Crm1 N-end rule degron strain (Fig EV4D) was generated by integrating pMK632 (Gnanasundram & Kos, 2015) into the *CRM1* locus in the presence of the pCM188 TET activator plasmid to create RSY2348 (Ubi-Leu-3HA-*CRM1*::*NATMX*) as described in detail in (Willis *et al*., 2020). In accordance with the Mediator nomenclature unification efforts (Bourbon *et al*, 2004) members of the Cdk8 module, the cyclin C (*SSN8*/*UME3*/*SRB11*), Cdk8 (*SSN3*/*UME5*/*SRB10*), *MED12* (*SSN5*) and (*MED13/UME2/SRB9/SSN2*) will use *CNC1*, *CDK8, MED12* and *MED13* gene designations, respectively.

Plasmids used in this study are listed in Extended Data Table 3. The wild-type epitope-tagged plasmids Vph1-mCherry, Nab2-mCherry, Med13-HA and *MED13* Y2H *GAL4* activating domain plasmids have been previously described (Stieg *et al*., 2018; Willis *et al*., 2020). The *GAL4-BD-SNX4* fusion plasmids were constructed by amplifying *SNX4* alleles from wild-type genomic DNA with oligonucleotides containing *Sal*I flanking sites and cloning into the *Sal*I site of the *GAL4* binding domain plasmid pAS2 (Wang & Solomon, 2012). The Crn1-GFP-Med13 plasmid was created by amplifying the first 400 amino acids of Crn1 from genomic DNA and cloning it in-frame to the N terminus of GFP-Med13 (pSW218). Plasmid construction details are available upon request. All constructs were verified by sequencing.

### Cell growth

Yeast cells were grown in either rich, non-selective medium (YPDA: 2% (w/v) glucose, 2% (w/v) Bacto peptone, 1% (w/v) yeast extract, 0.001% (w/v) adenine sulfate) or synthetic minimal dextrose medium (SD: 0.17% (w/v) yeast nitrogen base without amino acids and ammonium sulfate, 0.5% (w/v) ammonium sulfate, 1 x supplement mixture of amino acids, 2% (w/v) glucose) allowing plasmids selection as previously described (Cooper *et al*., 1997). For nitrogen starvation experiments, cells were grown as described (Journo *et al*, 2009).

### Cellular assays

RT-PCR analysis was executed as previously described in (Cooper *et al*., 2012; Willis *et al*., 2020). Oligonucleotides used during these studies are available upon request. All studies were conducted with three biological samples in technical triplicates. The standard deviation from three replicate reactions is indicated in the figures. P values were determined using the unpaired Students t-test. The six-day nitrogen starvation viability assays were executed exactly as described (Willis *et al*., 2020) with 30,000 cells counted per timepoint using FACS and the studies were conducted in biological duplicates. Data are mean ± standard deviation

### Western blot assays

Protein extracts for Western blot studies were prepared using a NaOH lysis procedure exactly as described in ((Willis *et al*., 2018). In short, protein extracts were prepared from 25 ml per timepoint with the exception of Med13-9xmyc cultures in which 50 ml was needed to visualize the protein. Proteins were separated on 6-10% SDS polyacrylamide gels depending upon their size using the Bio-RAD Mini-Trans Blot cell. To detect proteins, 1:5000 dilutions of anti-myc (UpState New York), anti-HA (Abcam) or anti-Pgk1 (Invitrogen) antibodies were used. Western blot signals were detected using 1:5000 dilutions of either goat anti-mouse or goat anti-rabbit secondary antibodies conjugated to alkaline phosphatase (Abcam) and CDP-Star chemiluminescence kit (Invitrogen, cat.#T2307). Signals were quantified relative to Pgk1 or Tubulin (Fig EV1C) controls using CCD camera imaging (Kodak Inc.) All degradation assays were performed in triplicate. Standard deviation and significance were calculated from the mean ± standard deviation using GraphPad Prism 7.

### Cleavage assays

Strains harboring GFP-fusion proteins were grown to mid-log in SD, washed and resuspended in SD-N. Protein extracts were prepared using NaOH as described above (25 mLs /TP). Proteins were separated using Invitrogen Blot^TM^ 4-12% Bis-Tris Plus gradient gels with 1X MOPS SDS running buffer (cat.#NW04122BOX). Proteins were transferred to PVDF membranes in 1X Blot^TM^ transfer buffer for 1h 30mins (cat.#BT00061). GFP tagged proteins were detected using 1:5000 dilution of Anti-GFP (WAKO) antibodies and goat anti-mouse secondary antibodies conjugated to alkaline phosphatase.

### Co-Immunoprecipitation

For co-immunoprecipitation experiments, 1 L of cells were grown to mid-log, washed and resuspended in SD-N media (250 mLs/TP). Protein extracts were prepared using a glass bead lysis method exactly as described in (Stieg *et al*., 2018), except Protein A beads were pre-washed with IP wash solution (500mM NaCl, 25mM Tris). 1 mg of total protein was immunoprecipitated per timepoint. Anti-GFP antibodies (Invitrogen) or Anti-HA antibodies (Abcam) were used for immunoprecipitations. Co-immunoprecipitation blot was probed with antibodies against HA (Abcam) or T7 (Novagen) epitope. Due to the drastic difference in size between Med13-3xHA and GFP tagged proteins (Snx4 and Atg17) input controls were run on separate gels. For Atg17 experiments endogenous Atg17-GFP was immunoprecipitated from whole lysates to obtain input controls. For all other input controls 50 μg of protein was resuspended in 2 x SDS-PAGE and separated on either 6% (Med13-3HA) or 10% (GFP-Snx4 and Atg17-GFP) SDS polyacrylamide gels. All co-immunoprecipitation experiments were performed in *pep4Δ prb1Δ.1* strains. (Stieg *et al*., 2018; Willis *et al*., 2018).

### Fluorescence Microscopy

For all microscopy experiments, cells were grown to mid-log phase, washed and resuspended in SD-N for the time points indicated. Deconvolved images were obtained using a Nikon microscope (Model E800) with a 100x objective with 1.2x camera magnification (Plan Fluor Oil, NA 1.3) and a CCD camera (Hamamatsu Model C4742). Data were collected using NIS software and processed using Image Pro software. All images of individual cells were optically sectioned (0.2 μM slices at 0.3 μM spacing) and deconvolved, and slices were collapsed to visualize the entire fluorescent signal within the cell. The nuclei were visualized in live cells using Hoechst staining (Cayman Chemical 15547). Hoechst (5 μM), dissolved in water, was added to cells growing in either SD or SD-N (5 μM) 30 min before they were visualized by microscopy. The vacuole was visualized in live cells in (Fig EV2A) using FM4-64 (Invitrogen, T3166) and phenylmethane-sulfonyl-fluoride (PMSF, Sigma P7626) treatment of cells was executed exactly as described (Journo *et al*, 2008). In order to optimize the visualization of the lowly expressed endogenous Med13-mNeongreen the Keyence microscope was used. This scope has a high sensitivity CCD and high-speed autofocus, low photobleaching mode that aids in monitoring Med13 localization in live cells. Deconvolution and processing capabilities are very limited with the compatible analyzer software. Single plane images were obtained using a Keyence BZ-X710 fluorescence microscope with a 100x objective with 1.0x camera magnification (PlanApoλ Oil, NA 1.45) and a CCD camera. Data were collected using BZ-X Analyzer software. Quantification of Med13-mNeongreen fluorescence within the vacuole was obtained using the Hybrid cell count function within the analyzer software (300 cells were counted per sample). For analysis single extraction settings were used. Red (vacuole, Vph1-mCherry) was set as the target area and green (Med13-mNeongreen) was set as the extraction area. The percentage of cells with vacuolar Med13-mNeongreen was calculated using Area ratio (1^st^) (ratio of the total area of the extracted areas to the target area) and cell count values. Percentages represent a ratio of extraction area to the target area.

### Statistics

All representative results included at least two independent biological experiments. P values were generated from Prism-GraphPad using unpaired Student’s t-tests; NS P ≥ 0.05; *P ≤ 0.05, **P ≤ 0.005; ***P ≤ 0.001; ****P ≤ 0.0001. All error bars indicate mean ± SD. For quantification of Med13-9myc degradation kinetics band intensities of each time point was first divided by unstressed, T=0 band intensity. These values were then divided by Pgk1 loading band intensity values which were also normalized to their T=0 intensities. P-values shown are relative to wild-type T=4 timepoints.

## ACKNOWLEDGMENTS

We thank C. De Virgilio, P. Herman and K. Cunningham for strains and plasmids. We especially thank the members of the R. Strich laboratory and the Cooper laboratory for critical reading of this manuscript.

## FUNDING

This work was supported by a grant from the National Institutes of Health awarded to K.F.C. (GM113196).

## COMPETING INTERESTS

The authors declare no competing or financial interests.

## AUTHOR CONTRIBUTIONS

SEH, SDW and KFC conceived the study and designed experiments. SEH and SDW performed experiments. KFC performed most of the microscopy experiments executed with the Nikon microscope. SEH and KFC wrote the manuscript.

## Supplementary Information

**Supplemental Table I.** Candidate proteins that interact with Med13 after 90 min 200mM rapamycin treatment.

| Select proteins obtained from Mass Spectroscopy where:<br>Counts were greater than 2-fold after 90 minutes of 200 ng/mL Rapamycin |  |  | Counts |  |
| --- | --- | --- | --- | --- |
| Systematic Name | Gene | Protein Function | T=0 | T=90 |
| **Other** |  |  |  |  |
| YER155C | *BEM2* | RhoGAP | 47 | 203 |
| YJL005W | *CYR1* | Adenylate cyclase | 19 | 50 |
| YHR016C | *YSC84* | Actin-binding protein | 7 | 18 |
| YNL271C | *BNI1* | Formin | 5 | 13 |
| YBR140C | *IRA1* | GTPase activator | 1 | 12 |
| YDR103W | *STE5* | MAPK scaffold | 4 | 12 |
| YGR032W | *GSC2* | spore wall assembly | 1 | 9 |
| YDL198C | *GGC1* | Mitochondrial GTP/GDP transporter | 3 | 7 |
| **Nuclear Activity** |  |  |  |  |
| YPR018W | *RLF2* | Chromatin assembly | 21 | 44 |
| YPL155C | *KIP2* | Kinesin | 6 | 34 |
| YDL140C | *RPO21* | RNA Polymerase II Subunit | 13 | 33 |
| YML065W | *ORC1* | DNA replication | 14 | 32 |
| YNL278W | *CAF120* | Transcriptional regulation | 7 | 27 |
| YDR216W | *ADR1* | Transcription factor | 12 | 26 |
| YER040W | *GLN3* | Transcriptional activator | 10 | 23 |
| YDR364C | *CDC40* | mRNA Splicing | 7 | 15 |
| YGL045W | *RIM8* | Activates Rim101 | 6 | 13 |
| YER088C | *DOT6* | DNA Binding Protein | 3 | 12 |
| YDL089W | *NUR1* | Mitotic Exit/Nuclear Periphery | 3 | 9 |
| YDL207W | *GLE1* | Cytoplasmic nucleoporin | 3 | 9 |
| YDL220C | *CDC13* | Telomere capping | 4 | 9 |
| ***Vacuole*** |  |  |  |  |
| YJL172W | *CPS1* | Vacuolar carboxypeptidase | 4 | 18 |
| YPR036W | *VMA13* | V-ATPase | 4 | 12 |
| YDR128W | *MTC5* | Vacuole/TORC1 | 3 | 8 |
| YLL048C | *YBT1* | Vacuole Fusion | 2 | 8 |
| ***Kinase/Phosphatase*** |  |  |  |  |
| YDR283C | *GCN2* | Kinase | 78 | 160 |
| YHR082C | *KSP1* | Kinase | 25 | 60 |
| YKL168C | *KKQ8* | Kinase | 13 | 36 |
| YDR122W | *KIN1* | Kinase | 4 | 28 |
| YOR039W | *CKB2* | Kinase | 8 | 26 |
| YDL025C | *RTK1* | Kinase | 3 | 18 |
| YOR061W | *CKA2* | Kinase | 5 | 15 |
| YGL019W | *CKB1* | Kinase | 4 | 13 |
| YMR165C | *PAH1* | Phosphatase | 4 | 13 |
| YIL035C | *CKA1* | Kinase | 5 | 12 |
| YAR014C | *BUD14* | Protein phosphatase regulator | 3 | 7 |
| ***Anabolism/Catabolism*** |  |  |  |  |
| YLR044C | *PDC1* | Pyruvate decarboxylase | 57 | 200 |
| YKL182W | *FAS1* | Fatty acid synthetase | 35 | 94 |
| YKL060C | *FBA1* | Glycolysis | 26 | 62 |
| YEL071W | *DLD3* | Lactate Dehydrogenase | 16 | 40 |
| YBR196C | *PGI1* | Phosphoglucose isomerase | 3 | 29 |
| YLR304C | *ACO1* | Aconitase | 10 | 25 |
| YLR153C | *ACS2* | Acetyl-coA synthetase | 8 | 24 |
| YOR136W | *IDH2* | Isocitrate dehydrogenase complex | 7 | 22 |
| YBR208C | *DUR1,2* | Urea amidolyase | 0 | 21 |
| YLR355C | *ILV5* | Amino acid biosynthesis | 1 | 19 |
| YGL009C | *LEU1* | Leucine biosynthesis | 7 | 17 |
| YKL218C | *SRY1* | Amino Acid Catabolism | 2 | 15 |
| YGL245W | *GUS1* | tRNA synthetase | 2 | 15 |
| YER024W | *YAT2* | Acetyltransferase/mitochondria | 1 | 11 |
| YBR221C | *PDB1* | Pyruvate Dehydrogenase | 3 | 10 |
| YDR341C | YDR341C | Arginyl-tRNA synthetase/mitochondria | 1 | 9 |
| YIL128W | *MET18* | Methionine biosynthesis/CIA Complex | 2 | 9 |
| YGL001C | *ERG26* | Ergosterol biosynthesis | 3 | 7 |
| YJL060W | *BNA3* | Aminotransferase | 3 | 7 |
| **Protein Trafficking** |  |  |  |  |
| YNL287W | *SEC21* | COPI vesicle coat | 9 | 23 |
| YBR102C | *EXO84* | ExoCytosis/Splicesome Assembly | 3 | 19 |
| YBL037W | *APL3* | Intracellular protein transport | 5 | 13 |
| YNL049C | *SFB2* | COPII vesicle coat | 5 | 12 |
| YNL246W | *VPS75* | Histone chaperone | 4 | 12 |
| YHR103W | *SBE22* | Golgi /Bud Growth | 2 | 11 |
| YLR114C | *AVL9* | Exocytic transport | 3 | 9 |
| YDR483W | *KRE2* | Golgi mannosyltransferase | 2 | 8 |
| YJL036W | *SNX4* | Sorting Nexin/CVT | 2 | 8 |
| YDR171W | *HSP42* | Chaperone | 3 | 7 |
| YDR186C | *SND1* | ER Targeting | 3 | 7 |
| **Ribosome** |  |  |  |  |
| YPL009C | YPL009C | Ribosome quality control complex | 14 | 63 |
| YKL014C | *URB1* | Ribosome biogenesis | 20 | 47 |
| YGR162W | *TIF4631* | Translation initiation factor | 12 | 37 |
| YDR333C | *RQC1* | Ribosome Quality Control | 13 | 31 |
| YMR049C | *ERB1* | Ribosome Biogenesis | 13 | 28 |
| YJR041C | *URB2* | Ribosome biogenesis | 1 | 11 |
| YNL224C | *SQS1* | Ribosome Biogenesis | 4 | 9 |
| **Unknown Function** |  |  |  |  |
| YPL032C | *SVL3* | Unknown Function/Vacuole | 16 | 59 |
| YER047C | *SAP1* | Unknown/AAA ATPase | 21 | 53 |
| YGR237C | YGR237C | Unknown Function | 11 | 28 |
| YLR187W | *SKG3* | Unknown Function | 9 | 19 |
| YLR219W | *MSC3* | Unknown Function | 8 | 18 |
| YBR007C | *DSF2* | Unknown/Bud Tip | 5 | 15 |
| YOR093C | YOR093C | Unknown Function | 6 | 15 |
| YBR225W | YBR225W | Unknown/Cell Wall | 5 | 13 |
| YDR251W | *PAM1* | Unknown Function | 2 | 13 |
| YGR196C | *FYV8* | Unknown Function | 4 | 11 |
| YDR239C | YDR239C | Unknown Function/Ribosome | 2 | 9 |
| ***Transporters/Permeases*** |  |  |  |  |
| YDR345C | *HXT3* | glucose transporter | 7 | 22 |
| YKR093W | *PTR2* | Peptide Transporter | 6 | 21 |
| YKR039W | *GAP1* | Amino Acid Permease | 4 | 20 |
| YJR152W | *DAL5* | Allantoate permease | 0 | 15 |
| YCL025C | *AGP1* | amino acid permease | 4 | 11 |
| ***Ubiquitin*** |  |  |  |  |
| YBL067C | *UBP13* | Ubiquitin Protease | 7 | 19 |
| YEL012W | *UBC8* | Ubiquitin Conjugating | 7 | 16 |
| YDL126C | *CDC48* | ATPase/Ubiquitin Binding | 6 | 15 |
| YDL122W | *UBP1* | Ubiquitin Protease | 3 | 11 |
| YDR328C | *SKP1* | Ubiquitin Ligase | 3 | 9 |
| YML088W | *UFO1* | F-box receptor protein | 2 | 8 |

**Supplemental Table II.**
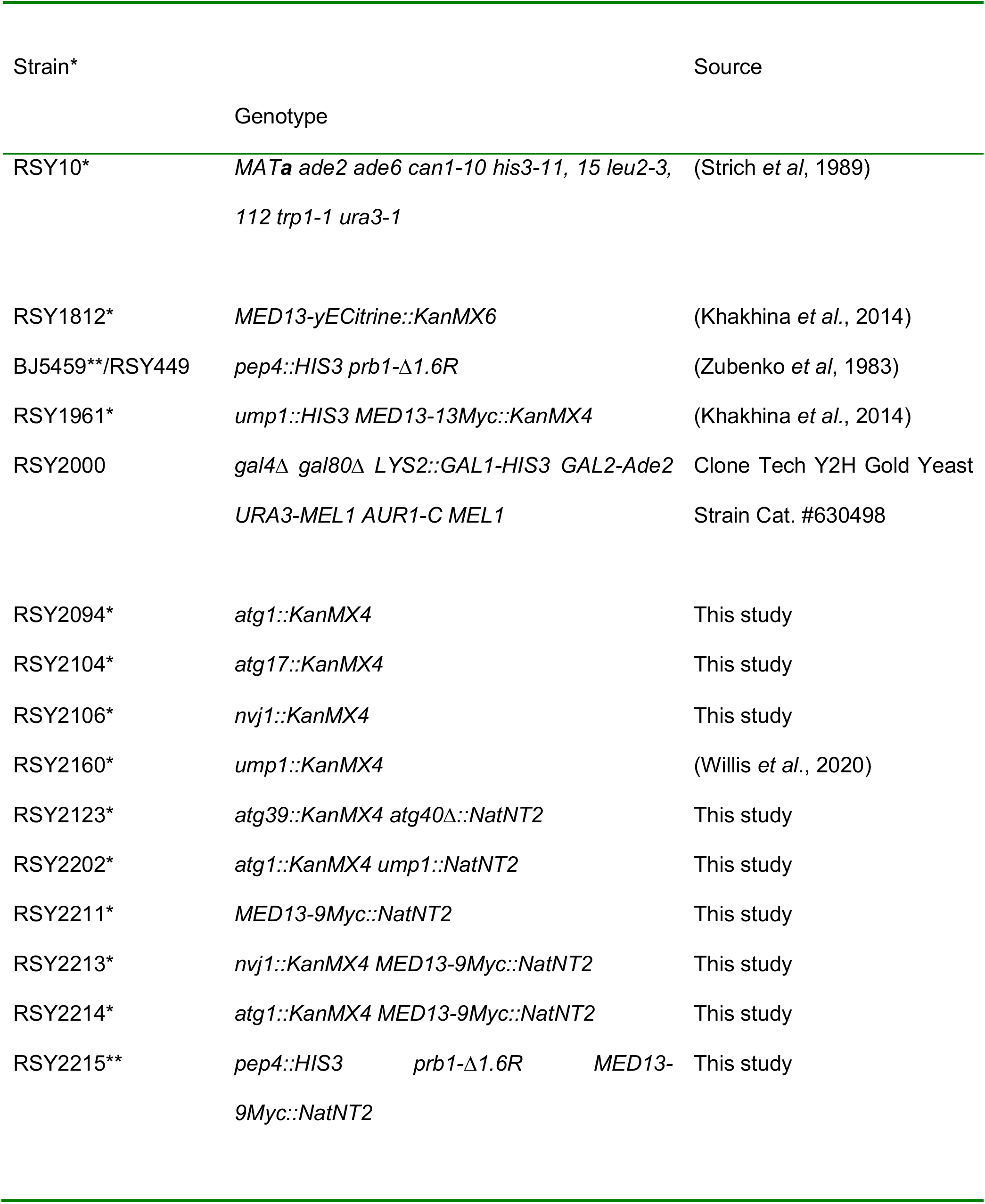
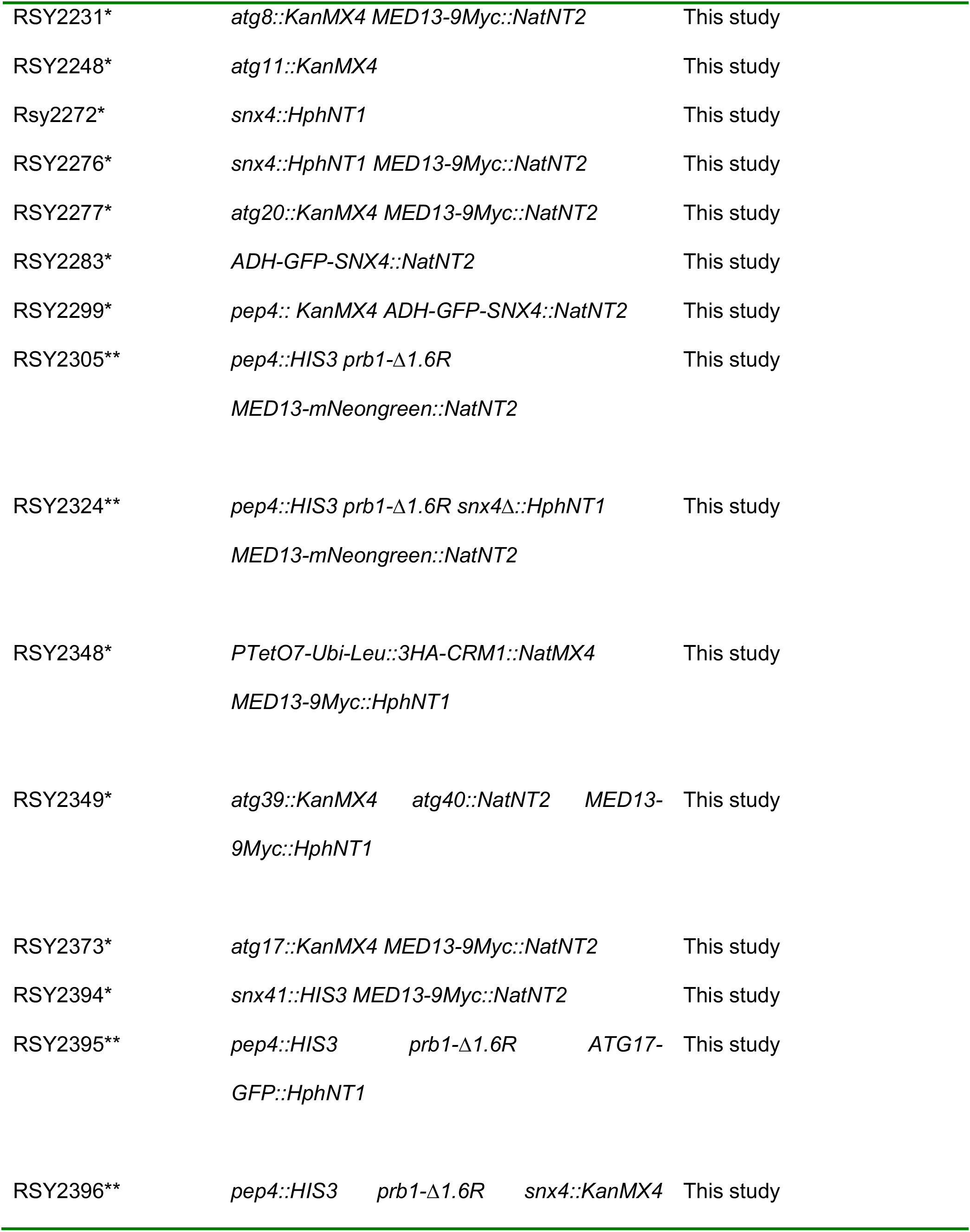
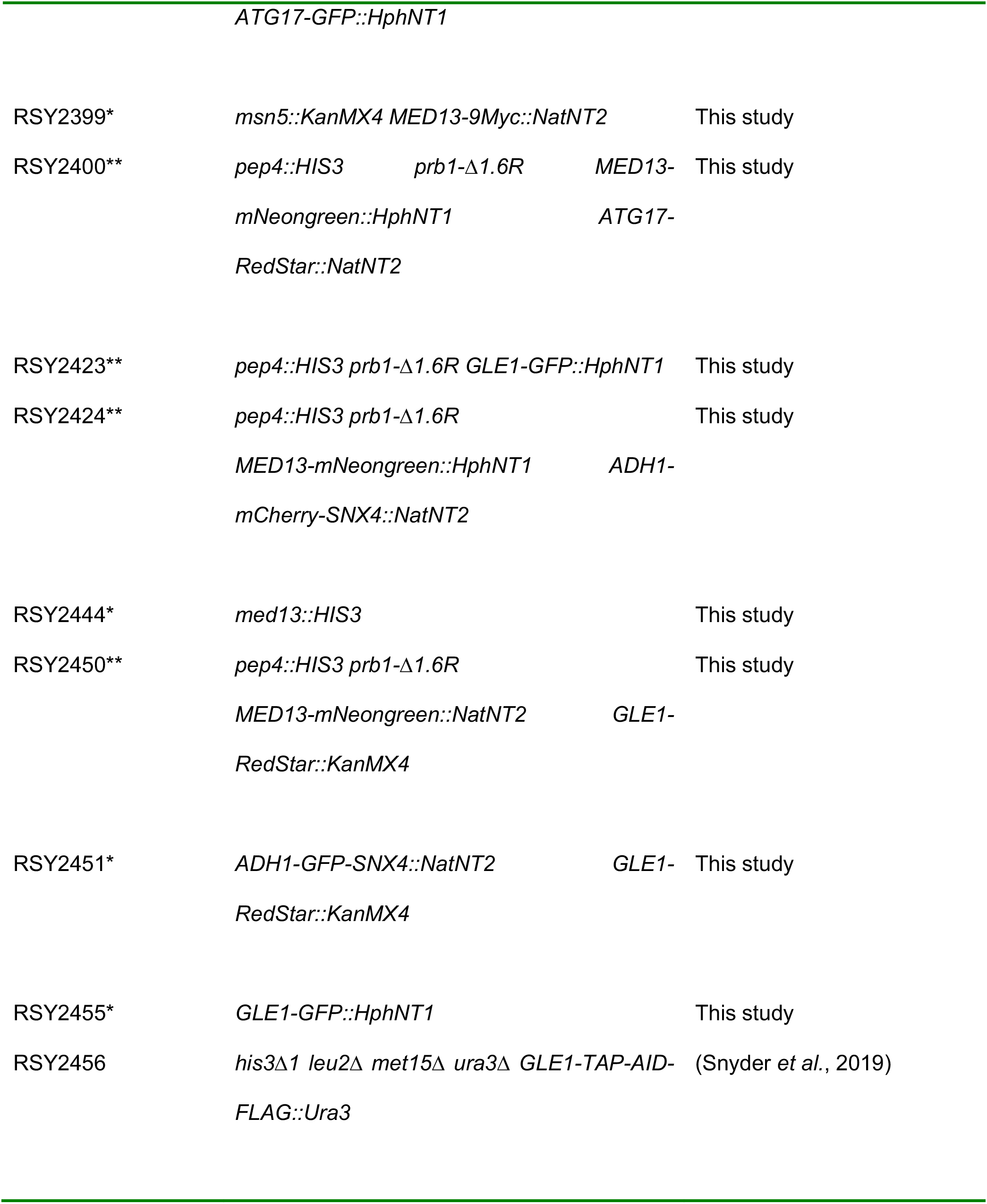
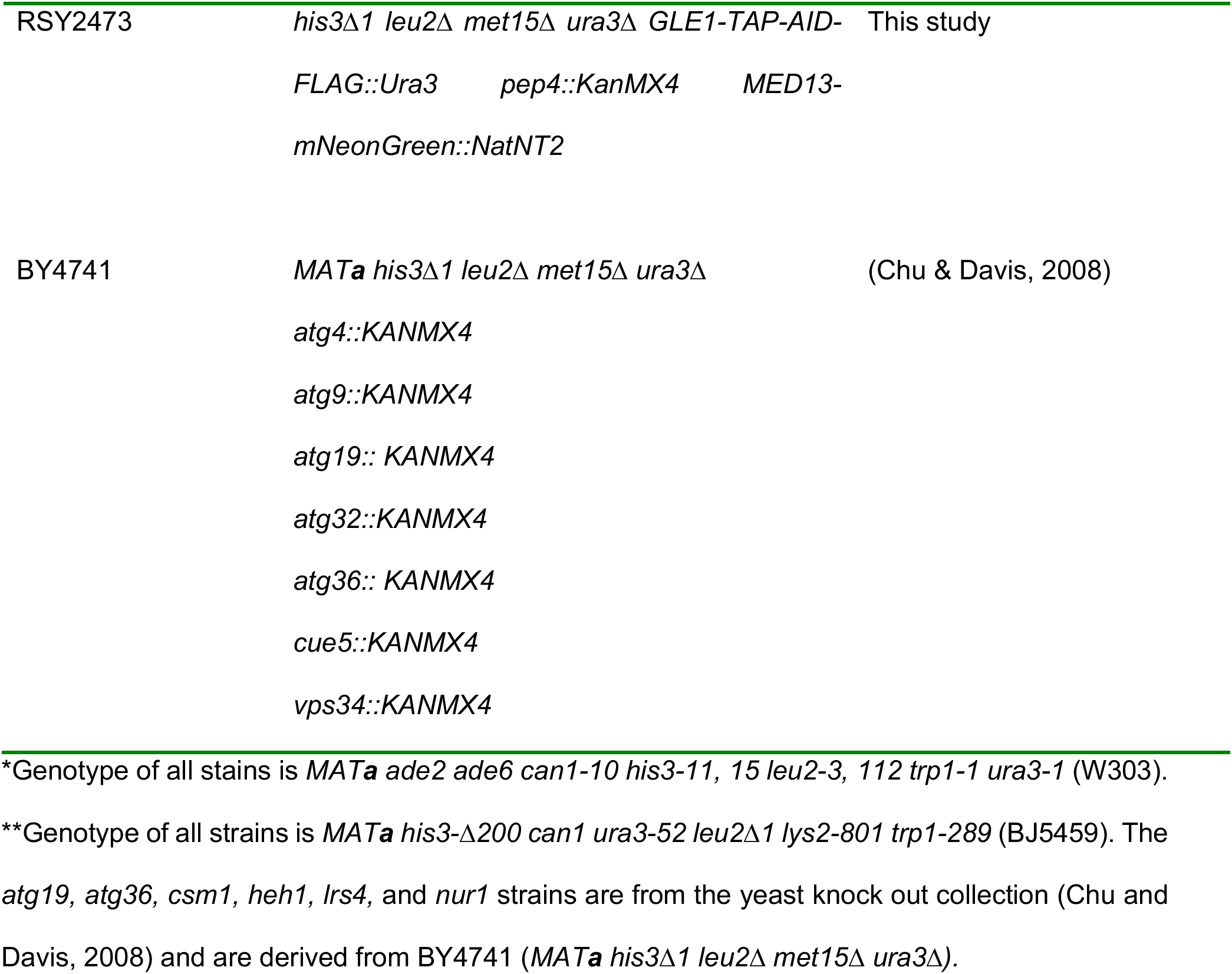
Yeast strains used in this study.

**Extended Data Table III.**
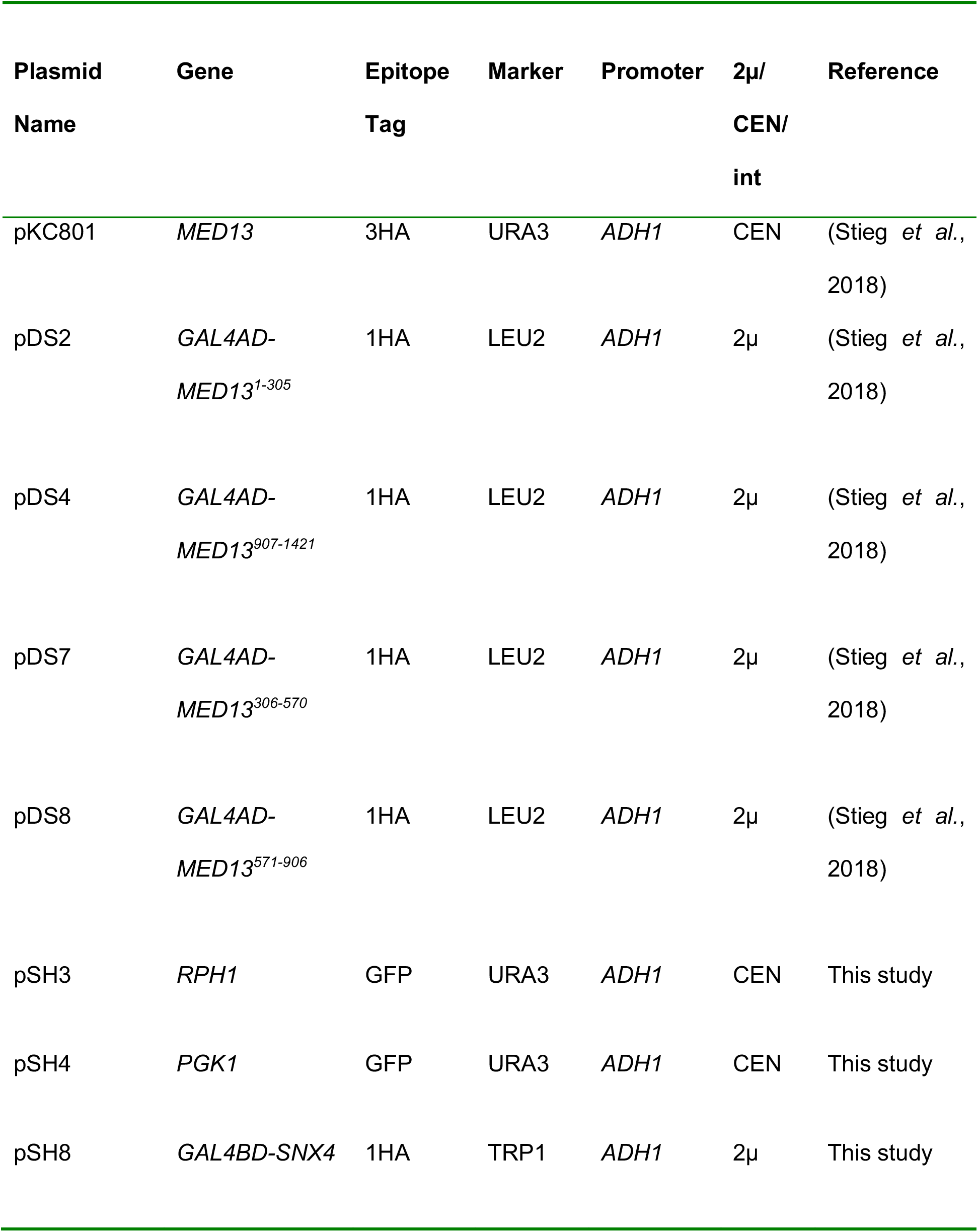
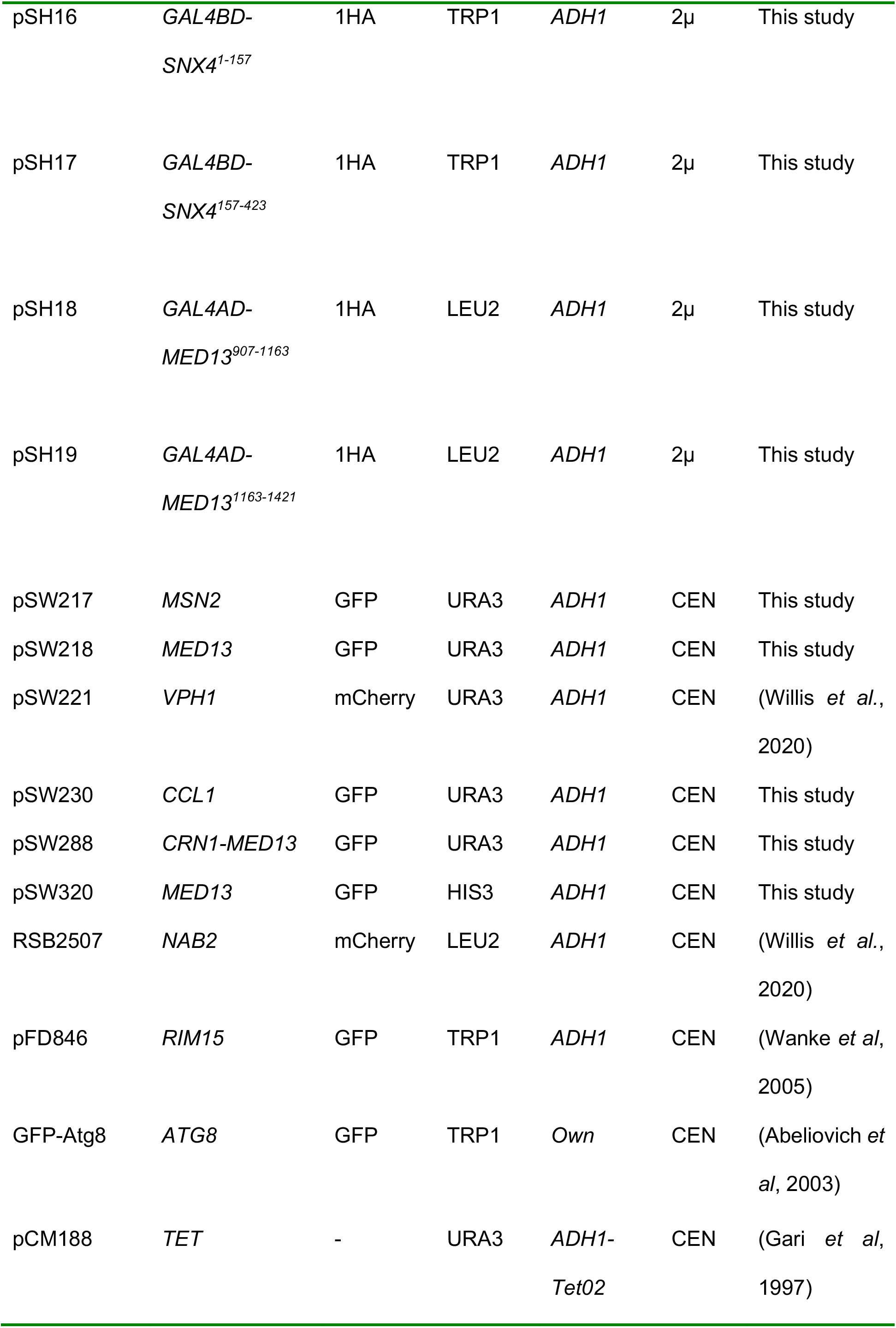
Plasmids used in this study.

| Plasmid Name | Gene | Epitope Tag | Marker | Promoter | 2μ/<br>CEN/<br>int | Reference |
| --- | --- | --- | --- | --- | --- | --- |
| pKC801 | *MED13* | 3HA | URA3 | *ADH1* | CEN | (Stieg *et al.*, 2018) |
| pDS2 | *GAL4AD-MED13*<sup>1-305</sup> | 1HA | LEU2 | *ADH1* | 2μ | (Stieg *et al.*, 2018) |
| pDS4 | *GAL4AD-MED13*<sup>907-1421</sup> | 1HA | LEU2 | *ADH1* | 2μ | (Stieg *et al.*, 2018) |
| pDS7 | *GAL4AD-MED13*<sup>306-570</sup> | 1HA | LEU2 | *ADH1* | 2μ | (Stieg *et al.*, 2018) |
| pDS8 | *GAL4AD-MED13*<sup>571-906</sup> | 1HA | LEU2 | *ADH1* | 2μ | (Stieg *et al.*, 2018) |
| pSH3 | *RPH1* | GFP | URA3 | *ADH1* | CEN | This study |
| pSH4 | *PGK1* | GFP | URA3 | *ADH1* | CEN | This study |
| pSH8 | *GAL4BD-SNX4* | 1HA | TRP1 | *ADH1* | 2μ | This study |
| pSH16 | *GAL4BD-SNX4<sup>1-157</sup>* | 1HA | TRP1 | *ADH1* | 2μ | This study |
| pSH17 | *GAL4BD-SNX4<sup>157-423</sup>* | 1HA | TRP1 | *ADH1* | 2μ | This study |
| pSH18 | *GAL4AD-MED13<sup>907-1163</sup>* | 1HA | LEU2 | *ADH1* | 2μ | This study |
| pSH19 | *GAL4AD-MED13<sup>1163-1421</sup>* | 1HA | LEU2 | *ADH1* | 2μ | This study |
| pSW217 | *MSN2* | GFP | URA3 | *ADH1* | CEN | This study |
| pSW218 | *MED13* | GFP | URA3 | *ADH1* | CEN | This study |
| pSW221 | *VPH1* | mCherry | URA3 | *ADH1* | CEN | (Willis *et al.*, 2020) |
| pSW230 | *CCL1* | GFP | URA3 | *ADH1* | CEN | This study |
| pSW288 | *CRN1-MED13* | GFP | URA3 | *ADH1* | CEN | This study |
| pSW320 | *MED13* | GFP | HIS3 | *ADH1* | CEN | This study |
| RSB2507 | *NAB2* | mCherry | LEU2 | *ADH1* | CEN | (Willis *et al.*, 2020) |
| pFD846 | *RIM15* | GFP | TRP1 | *ADH1* | CEN | (Wanke *et al.*, 2005) |
| GFP-Atg8 | *ATG8* | GFP | TRP1 | *Own* | CEN | (Abeliovich *et al.*, 2003) |
| pCM188 | *TET* | - | URA3 | *ADH1-* | CEN | (Gari et al, |
|  |  |  |  | *Tet02* |  | 1997) |

**Supplemental Figure 1.**
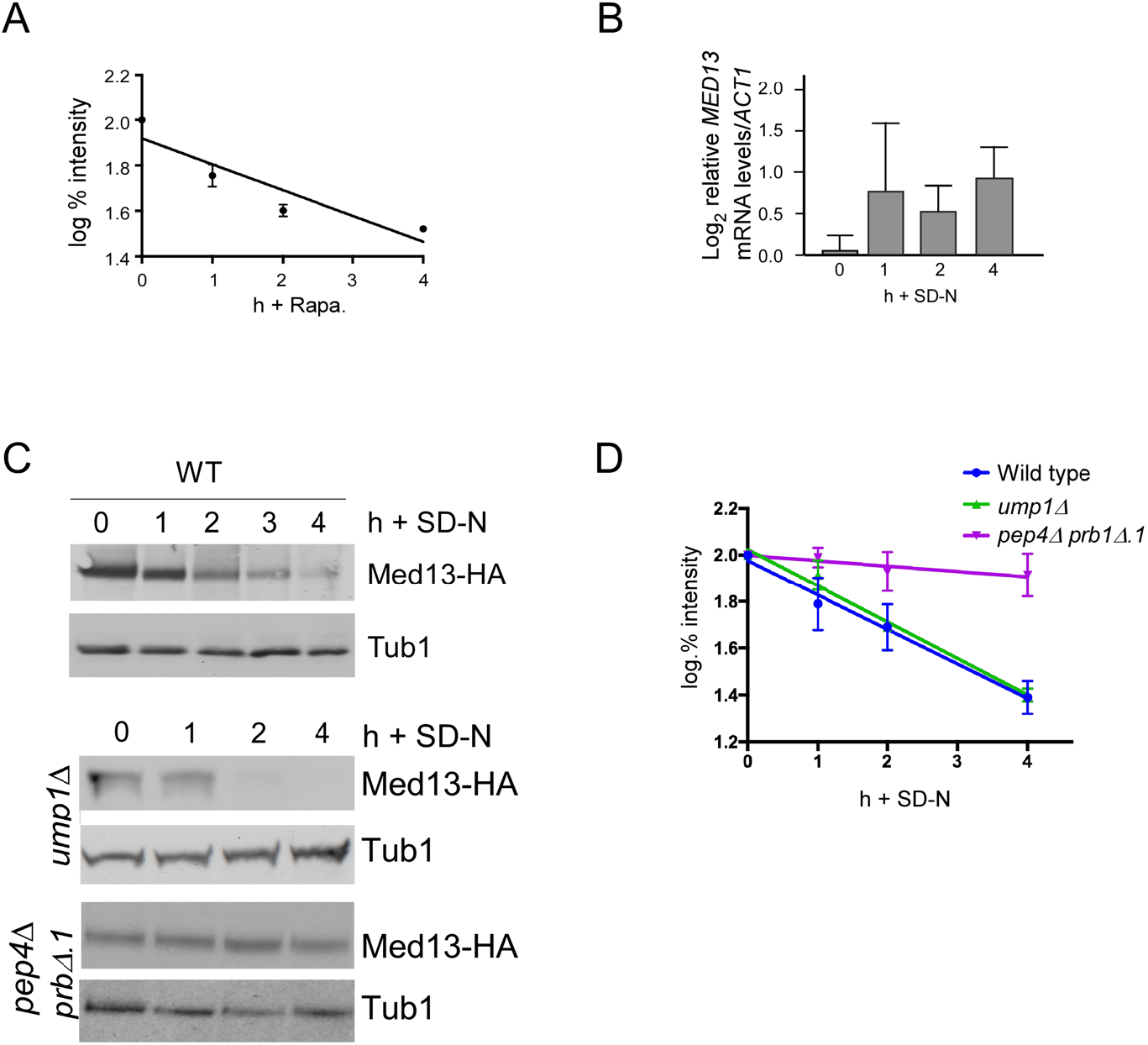
Med13 is degraded in response to rapamycin. **A** Degradation kinetics of endogenous Med13-9myc protein levels in wild-type cells treated with 200 ng/mL rapamycin. Error bars indicate S.D., N=3 biological samples. **B** RT-qPCR analysis probing for *MED13* mRNA expression in wild-type cells following nitrogen starvation. ΔΔCt results for relative fold change (log_2_) values using wild-type unstressed cells as a control. Transcript levels are given relative to the internal *ACT1* mRNA control. **C** Western blot analysis of Med13-3HA protein levels, expressed from a single copy functional plasmid (Stieg *et al*., 2018) (pKC801) in wild-type (RSY10), *ump1Δ* (RSY2160) and *pep4Δ prb1Δ.1* (RSY449) cells following nitrogen starvation. Pgk1 was used as a protein loading control in all experiments. **D** Quantification of Med13-3HA degradation kinetics in indicated mutants obtained from c. Error bars indicate S.D., N=3 biological samples.

**Supplemental Figure 2.**
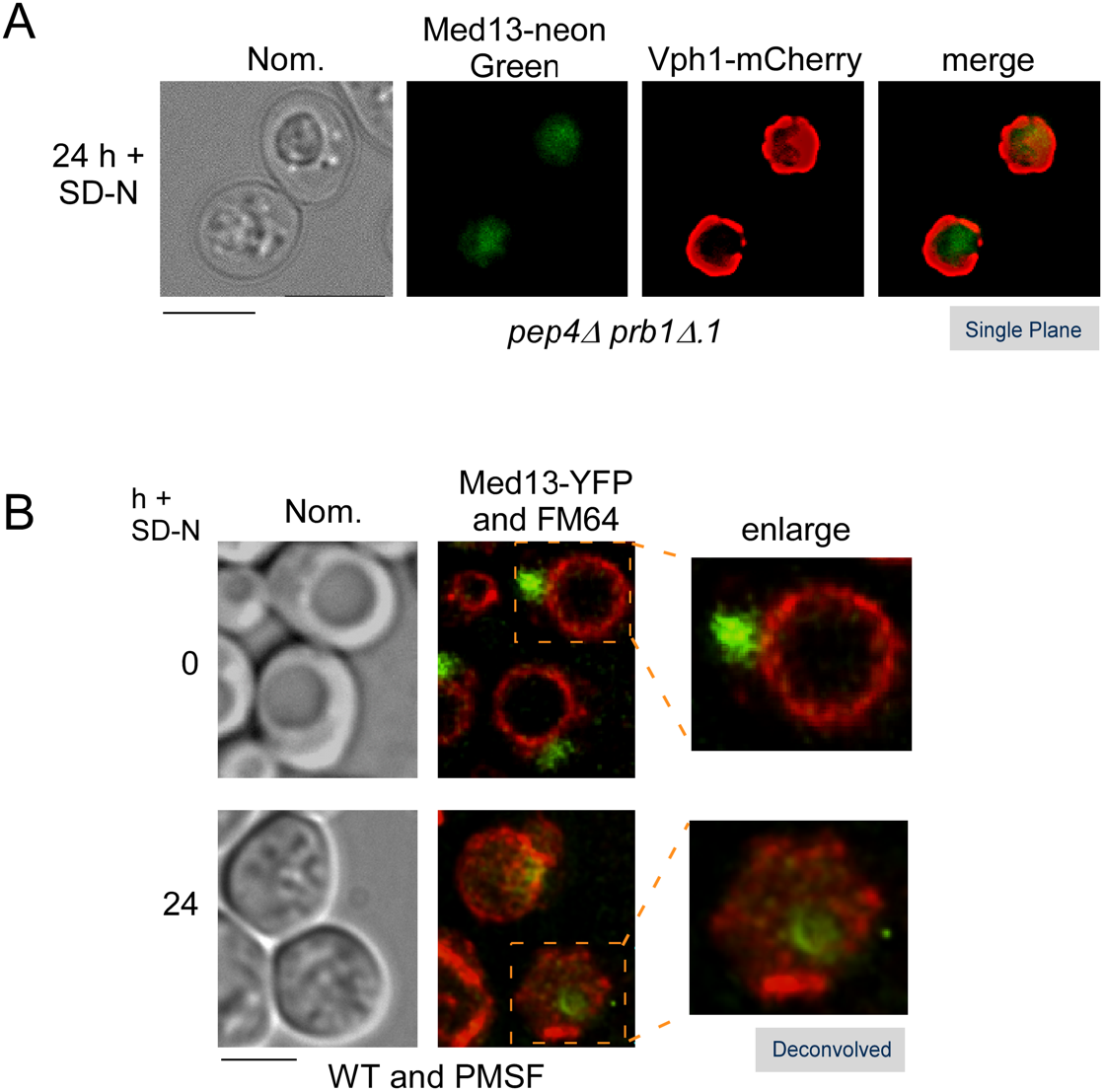
Med13 accumulates within the vacuole following nitrogen starvation. **A** Fluorescence microscopy of *pep4Δ prb1Δ.1* cells expressing endogenous Med13-mNeongreen and Vph1-mCherry (vacuole marker) after 24 h in SD-N. Representative single plane images are shown. Bar = 5μm. **B** Fluorescence microscopy of wild-type cells expressing endogenous Med13-YFP (RSY1812) treated with 200 mM PMSF before and after 24 h in SD-N media. FM4-64 staining was used to visualize the vacuole. Representative deconvolved images are shown. Bar = 5μm.

**Supplemental Figure 3.**
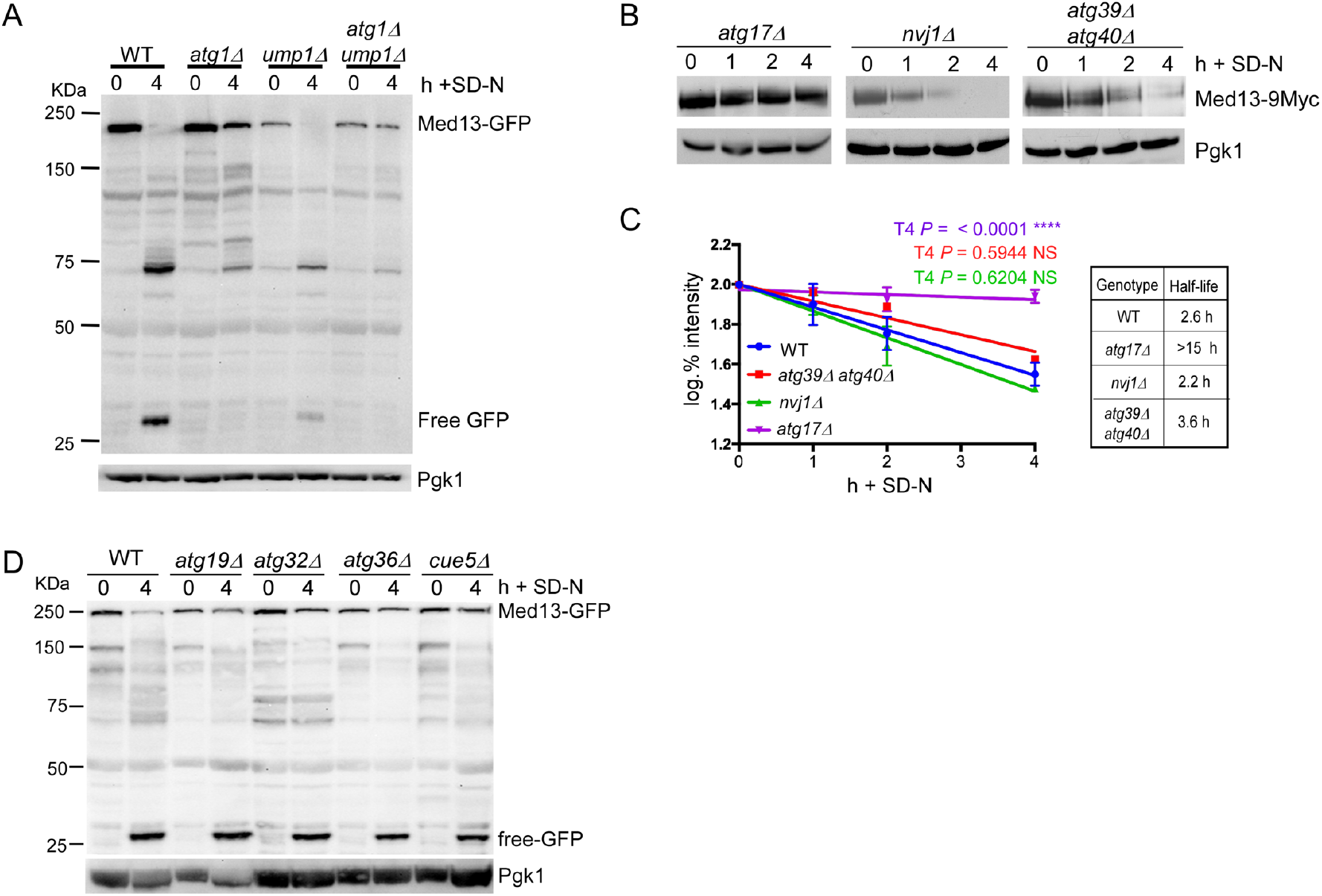
The autophagic degradation of Med13 does not require known selective autophagy adapter proteins. **A** Western blot analysis of Med13-GFP cleavage assays in WY (RSY10), *atg1Δ* (RSY2094), (RSY2160) and *atg1Δ ump1Δ* (RSY2202) cells after nitrogen starvation. **B** Western blot analysis of endogenous Med13-9xmyc degradation kinetics in *atg17Δ* cells as well as in strains defective in known nucleophagy pathways following nitrogen starvation. **C** Degradation kinetics and half-life of Med13 protein levels obtained in a. Error bars indicate S.D., N=3 biological samples. **D** Wild-type BY4741 cells and the indicated mutants expressing Med13-GFP (pSW218) were nitrogen-starved for the indicated times and the accumulation of free GFP was monitored by Western blot analysis using anti-GFP antibodies. Pgk1 levels were used as a protein loading control for all experiments. Degradation kinetics of Med13-GFP are slower in BY4741 cells which are more resistant to stress than the W303 strain. However, all these mutants accumulated free GFP which is indicative of vacuolar degradation.

**Supplemental Figure 4.**
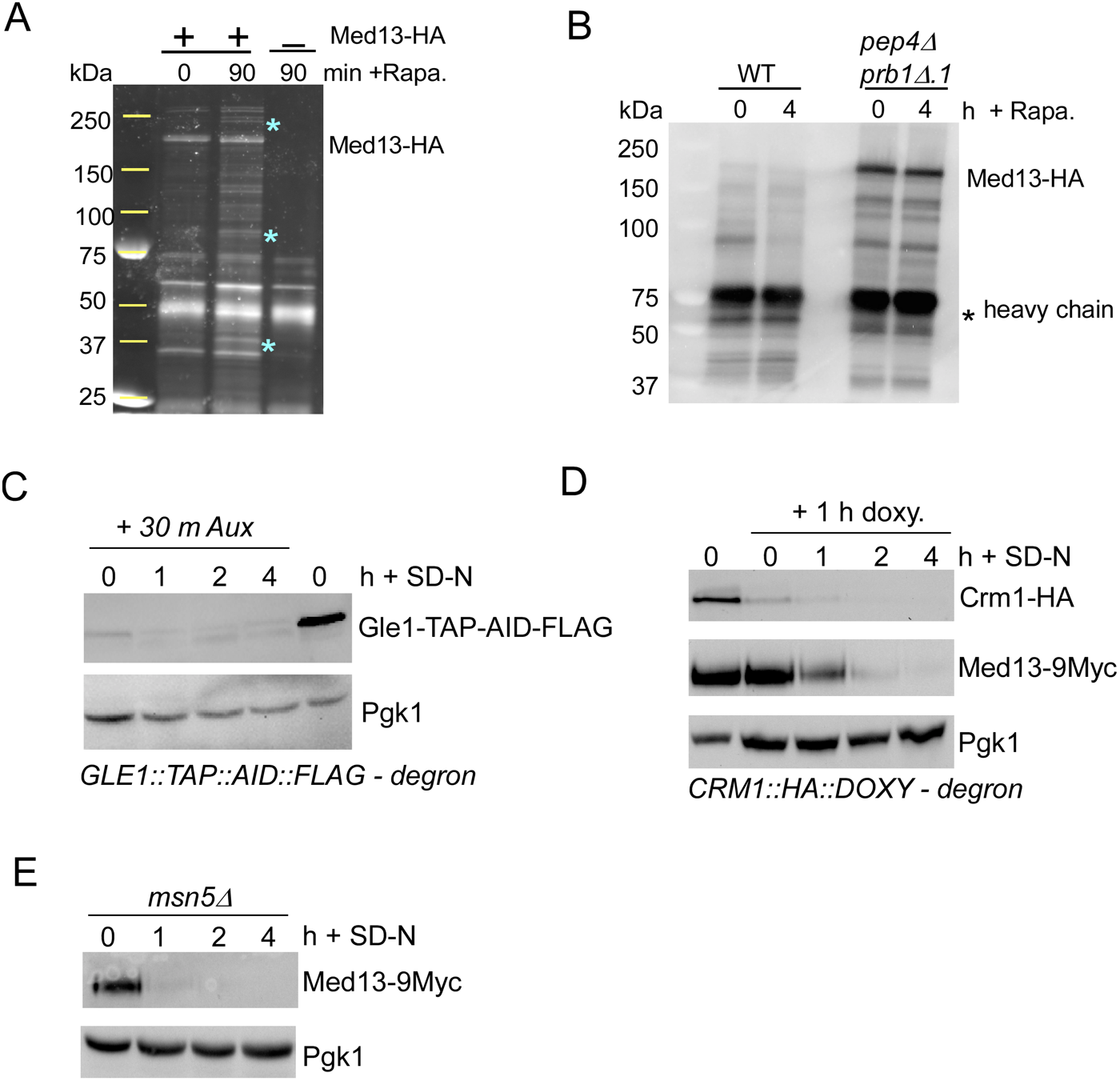
Mass spectrometry screen to identify proteins required for Med13 vacuolar degradation. **A** SDS-PAGE gel of immunoprecipitation assays with Med13 in unstressed cells or cells treated with 50nM rapamycin for 90 minutes. Spyro Ruby stain was used to visualize proteins immunoprecipitated with Med13-3xHA from whole cell lysate. Asterisks denote bands that were present in stressed samples and absent in unstressed samples. **B** Western blot analysis of immunoprecipitation experiments performed in wild-type and *pep4Δprb1Δ.1* cells. Cells harboring an expression plasmid for Med13-3HA were grown to mid-log and treated with 50nM rapamycin for 4 h. **C** Gle1 auxin inducible degron-degron system. The BY4741 strain harboring endogenous *GLE1-TAP-AID-FLAG* were grown to mid-log and treated with 250 μM Auxin. Cells were then washed and resuspended in SD-N media containing 250 μM Auxin for indicated time points. Gle1 protein depletion was monitored via Western blot analysis using antibodies against FLAG. **D** Crn1-doxycyclin inducible degron system. Cells expressing Crm1-HA under a tetracycline repressible promoter and endogenous Med13-9xmyc (RSY2348) were grown to mid-log. A sample was removed prior to the addition of doxycycline for Western blot analysis to visualize Crm1-HA (far left T=0 bands). The remaining culture was treated with doxycycline for 1 hour prior to switching cells to SD-N media with doxycycline. Western blot analysis was used to monitor Crm1-HA (top panel) and Med13-9xmyc (bottom panel) degradation following doxycycline and nitrogen starvation treatment. **E** Western blot analysis of Med13-9xmyc protein levels following nitrogen starvation in *msn5Δ* cells (RSY2399).

**Supplemental Figure 5.**
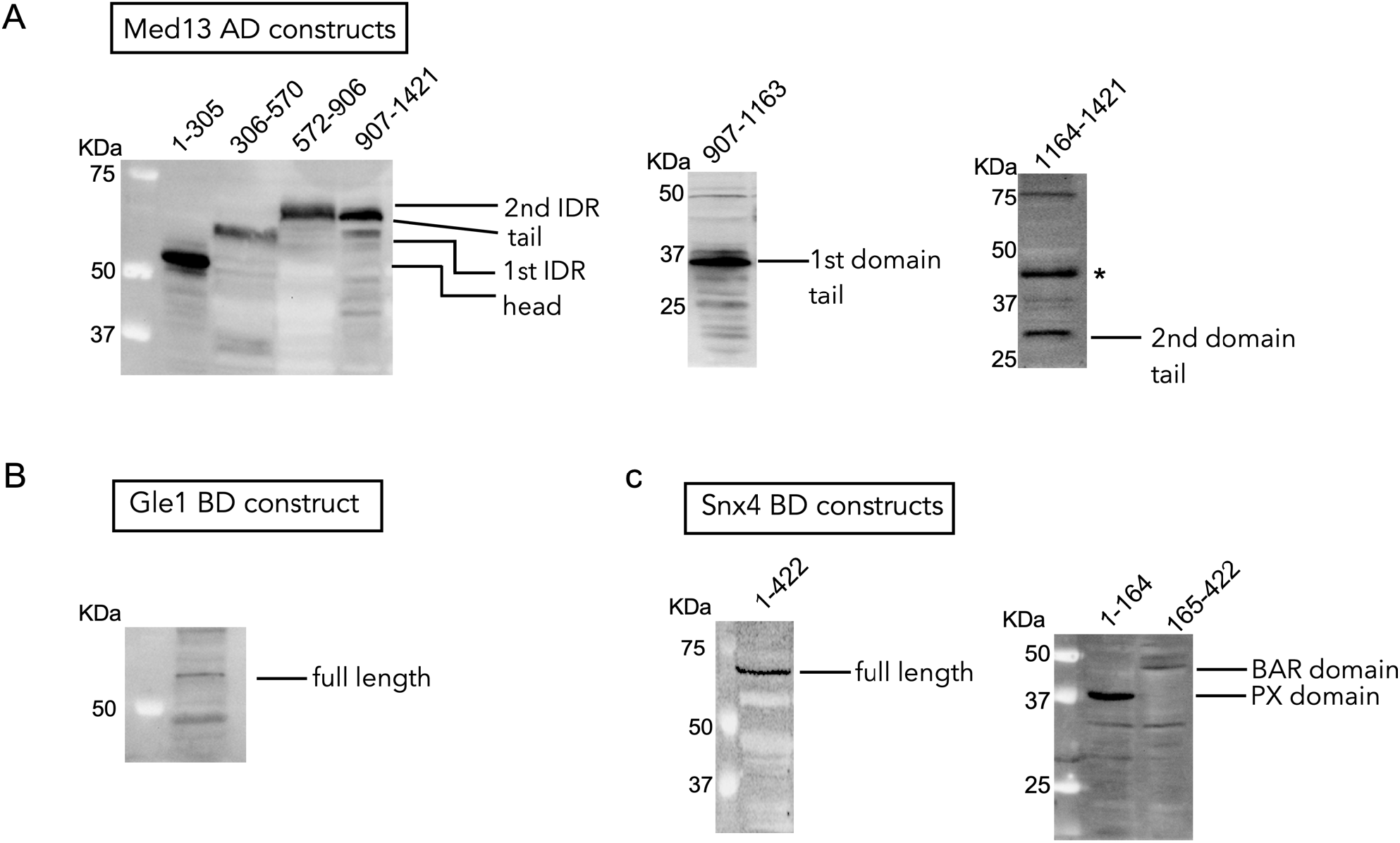
Western blot controls for Y2H plasmids. **A,** Western blot analysis of cells expressing indicated Med13 activating domain Y2H constructs. **B** and **C** Western blot analysis of cells expressing of Gle1 and Snx4 binding domain Y2H constructs respectively.

**Supplemental Figure 6.**
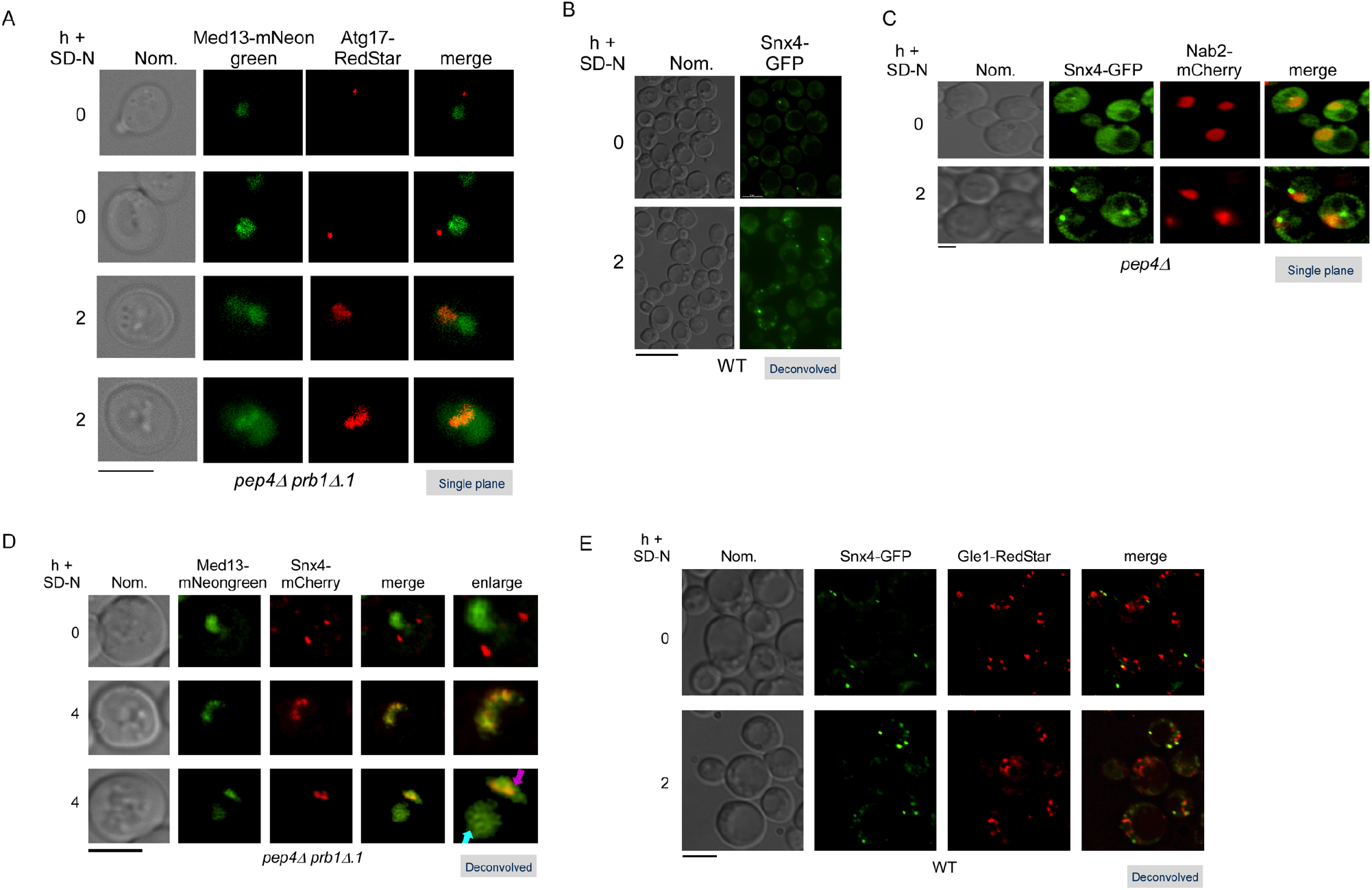
Snx4 localizes to the nuclear periphery following nitrogen starvation. **A** Fluorescence microscopy monitoring colocalization of endogenous Med13-mNeongreen and Atg17-Redstar after 2 h of nitrogen starvation in *pep4Δ prb1Δ.1* cells (RSY2400). Representative single plane images are shown. Bar = 5μm. **B** Fluorescence microscopy of GFP-Snx4 localization in wild-type cells following 2 h of nitrogen starvation. The number of foci increases following stress. Representative deconvolved images are shown. Bar = 5μm. **C** Fluorescence microscopy of *pep4Δ* cells expressing GFP-Snx4 and harboring Nab2-mCherry (nuclear marker) before and after nitrogen starvation. Representative deconvolved images are shown. Bar = 5μm.) **D** Fluorescence microscopy of endogenous Med13-mNeongreen and mCherry-Snx4 in *pep4Δ prb1Δ.1* cells before and after nitrogen starvation. Representative deconvolved images are shown. Vacuolar Med13 population indicated by the blue arrow and perinuclear Med13-Snx4 colocalization indicated by the pink arrow. Bar = 5μm. **E** Fluorescence microscopy of colocalization experiments performed in wild-type cells expressing GFP-Snx4 and endogenous Gle1-RedStar. Cells were grown to mid-log, washed and resuspended in SD-N media for 2 h. Representative large field view deconvolved images are shown. Bar = 5μm.

**Supplemental Figure 7.**
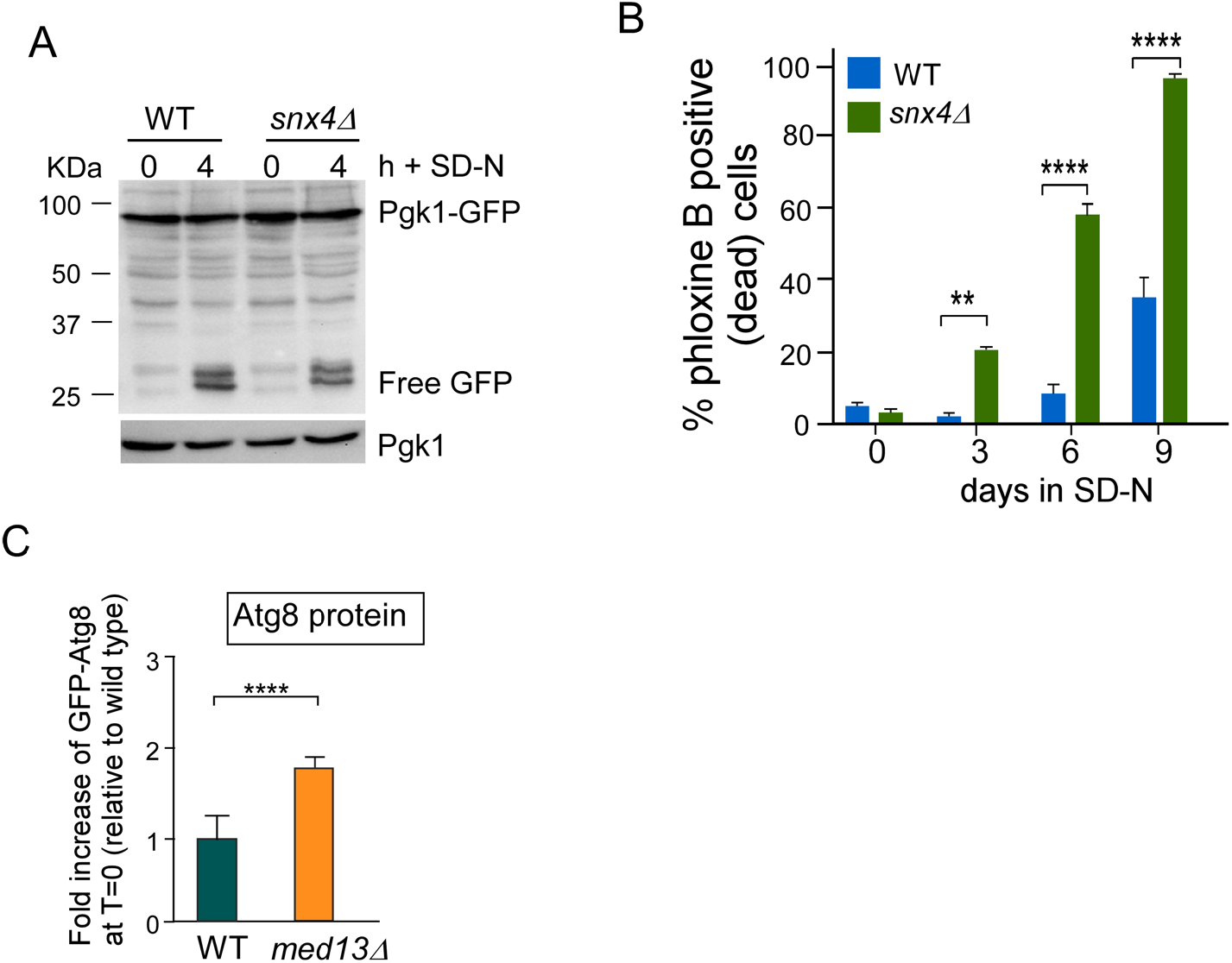
Snx4 is required for viability in long-term nitrogen starvation. **A** Western blot analysis of Pgk1-GFP (pSH4) cleavage assays in wild-type and *snx4Δ* cells following nitrogen starvation. Endogenous Pgk1 protein levels were used as a loading control. **B** Wild-type and *snx4Δ* cells were grown to mid-log, washed and resuspended in SD-N media for the indicated number of days. The percentage of inviable cells within the population was determined using pholxine B staining and fluorescence activated cell analysis (FAC). Quantification of N=2 independent biological experiments. Data are mean ± S.D. *\*\* P* = 0.0012, *\*\*\*\* P* = <0.0001 (Willis *et al*., 2020) **C** Quantification Atg8 protein levels in wild-type and *med13Δ* cells relative to Pgk1 loading control. Quantification of N=3 biological samples. Data are mean ± S.D. *\*\*\*\* P* = <0.0001

**Source Data Figure 2.**
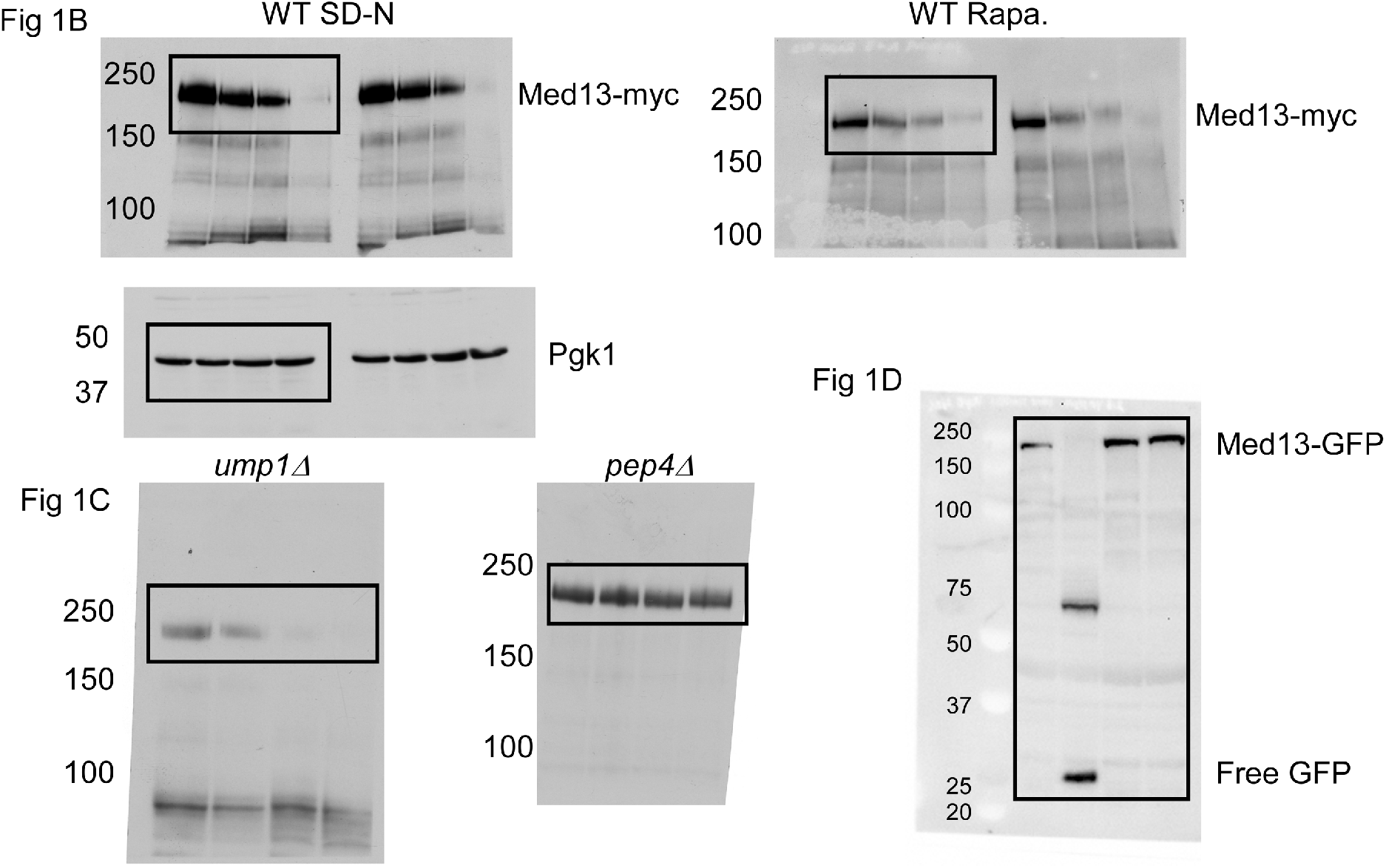
Source Data for Figure 1. Black boxes contain cropped representative images depicted in figure 1.

**Source Data Figure 2.**
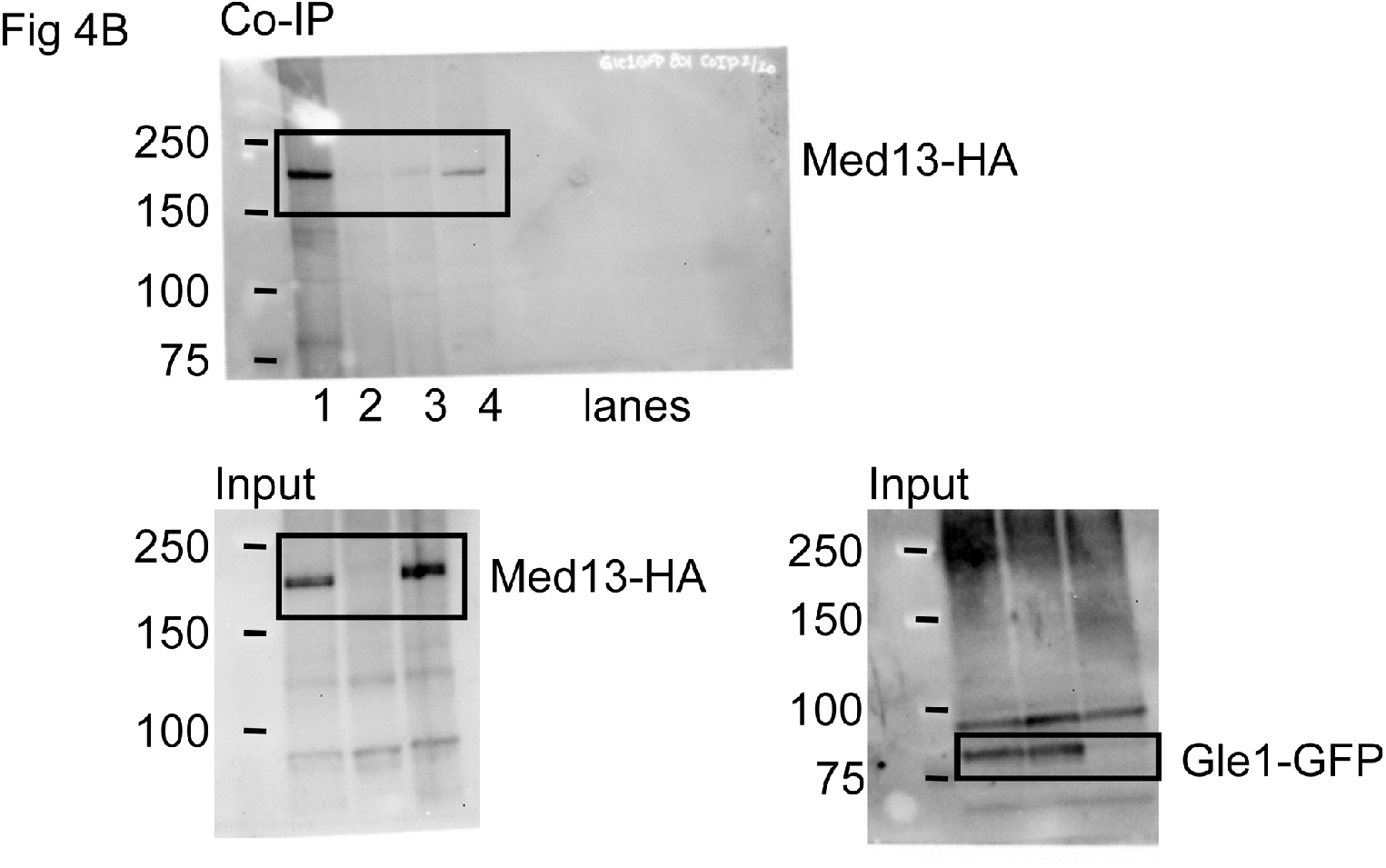
Source Data for Figure 4B. Black boxes contain representative cropped images depicted in figure 4.

## REFERENCES

Abeliovich H, Zhang C, Dunn WA, Jr., Shokat KM, Klionsky DJ (2003) Chemical genetic analysis of Apg1 reveals a non-kinase role in the induction of autophagy. Mol Biol Cell 14: 477–490

Akoulitchev S, Chuikov S, Reinberg D (2000) TFIIH is negatively regulated by cdk8-containing mediator complexes. Nature 407: 102–106.

Alcazar-Roman AR, Tran EJ, Guo S, Wente SR (2006) Inositol hexakisphosphate and Gle1 activate the DEAD-box protein Dbp5 for nuclear mRNA export. Nat Cell Biol 8: 711–716

Aryanpur PP, Regan CA, Collins JM, Mittelmeier TM, Renner DM, Vergara AM, Brown NP, Bolger TA (2017) Gle1 Regulates RNA Binding of the DEAD-Box Helicase Ded1 in Its Complex Role in Translation Initiation. Mol Cell Biol 37

Bartholomew CR, Suzuki T, Du Z, Backues SK, Jin M, Lynch-Day MA, Umekawa M, Kamath A, Zhao M, Xie Z et al (2012) Ume6 transcription factor is part of a signaling cascade that regulates autophagy. Proc Natl Acad Sci U S A 109: 11206–11210

Bean BD, Davey M, Conibear E (2017) Cargo selectivity of yeast sorting nexins. Traffic 18: 110–122

Bernard A, Jin M, Gonzalez-Rodriguez P, Fullgrabe J, Delorme-Axford E, Backues SK, Joseph B, Klionsky DJ (2015) Rph1/KDM4 mediates nutrient-limitation signaling that leads to the transcriptional induction of autophagy. Curr Biol 25: 546–555

Bourbon HM, Aguilera A, Ansari AZ, Asturias FJ, Berk AJ, Bjorklund S, Blackwell TK, Borggrefe T, Carey M, Carlson M et al (2004) A Unified Nomenclature for Protein Subunits of Mediator Complexes Linking Transcriptional Regulators to RNA Polymerase II. Mol Cell 14: 553–557

Chu AM, Davis RW (2008) High-throughput creation of a whole-genome collection of yeast knockout strains. Methods Mol Biol 416: 205–220

Cooper KF, Khakhina S, Kim SK, Strich R (2014) Stress-induced nuclear-to-cytoplasmic translocation of cyclin C promotes mitochondrial fission in yeast. Dev Cell 28: 161–173

Cooper KF, Mallory MJ, Smith JB, Strich R (1997) Stress and developmental regulation of the yeast C-type cyclin Ume3p (Srb11p/Ssn8p). EMBO J 16: 4665–4675

Cooper KF, Mallory MJ, Strich R (1999) Oxidative stress-induced destruction of the yeast C-type cyclin Ume3p requires phosphatidylinositol-specific phospholipase C and the 26S proteasome. Mol Cell Biol 19: 3338–3348

Cooper KF, Scarnati MS, Krasley E, Mallory MJ, Jin C, Law MJ, Strich R (2012) Oxidative-stress-induced nuclear to cytoplasmic relocalization is required for Not4-dependent cyclin C destruction. J Cell Sci 125: 1015–1026

Delorme-Axford E, Klionsky DJ (2018) Transcriptional and post-transcriptional regulation of autophagy in the yeast Saccharomyces cerevisiae. J Biol Chem

Deng Y, Qu Z, Naqvi NI (2013) The role of snx41-based pexophagy in magnaporthe development. PLoS One 8: e79128

Farre JC, Subramani S (2016) Mechanistic insights into selective autophagy pathways: lessons from yeast. Nat Rev Mol Cell Biol 17: 537–552

Fernandez-Martinez J, Kim SJ, Shi Y, Upla P, Pellarin R, Gagnon M, Chemmama IE, Wang J, Nudelman I, Zhang W et al (2016) Structure and Function of the Nuclear Pore Complex Cytoplasmic mRNA Export Platform. Cell 167: 1215–1228 e1225

Folkmann AW, Noble KN, Cole CN, Wente SR (2011) Dbp5, Gle1-IP6 and Nup159: a working model for mRNP export. Nucleus 2: 540–548

Fu N, Yang X, Chen L (2018) Nucleophagy Plays a Major Role in Human Diseases. Curr Drug Targets 19: 1767–1773

Ganesan V, Willis SD, Chang KT, Beluch S, Cooper KF, Strich R (2019) Cyclin C directly stimulates Drp1 GTP affinity to mediate stress-induced mitochondrial hyperfission. Mol Biol Cell 30: 302–311

Gari E, Piedrafita L, Aldea M, Herrero E (1997) A set of vectors with a tetracycline-regulatable promoter system for modulated gene expression in Saccharomyces cerevisiae. Yeast 13: 837–848

Gnanasundram SV, Kos M (2015) Fast protein-depletion system utilizing tetracycline repressible promoter and N-end rule in yeast. Mol Biol Cell 26: 762–768

Hettema EH, Lewis MJ, Black MW, Pelham HR (2003) Retromer and the sorting nexins Snx4/41/42 mediate distinct retrieval pathways from yeast endosomes. EMBO J 22: 548–557

Hollenstein DM, Gomez-Sanchez R, Ciftci A, Kriegenburg F, Mari M, Torggler R, Licheva M, Reggiori F, Kraft C (2019) Vac8 spatially confines autophagosome formation at the vacuole in S. cerevisiae. J Cell Sci 132

Hollenstein DM, Kraft C (2020) Autophagosomes are formed at a distinct cellular structure. Curr Opin Cell Biol 65: 50–57

Hu YB, Dammer EB, Ren RJ, Wang G (2015) The endosomal-lysosomal system: from acidification and cargo sorting to neurodegeneration. Transl Neurodegener 4: 18

Humphries CL, Balcer HI, D’Agostino JL, Winsor B, Drubin DG, Barnes G, Andrews BJ, Goode BL (2002) Direct regulation of Arp2/3 complex activity and function by the actin binding protein coronin. J Cell Biol 159: 993–1004

Hutten S, Kehlenbach RH (2007) CRM1-mediated nuclear export: to the pore and beyond. Trends Cell Biol 17: 193–201

Janke C, Magiera MM, Rathfelder N, Taxis C, Reber S, Maekawa H, Moreno-Borchart A, Doenges G, Schwob E, Schiebel E et al (2004) A versatile toolbox for PCR-based tagging of yeast genes: new fluorescent proteins, more markers and promoter substitution cassettes. Yeast 21: 947–962

Jeronimo C, Langelier MF, Bataille AR, Pascal JM, Pugh BF, Robert F (2016) Tail and Kinase Modules Differently Regulate Core Mediator Recruitment and Function In Vivo. Mol Cell 64: 455–466

Jeronimo C, Robert F (2017) The Mediator Complex: At the Nexus of RNA Polymerase II Transcription. Trends Cell Biol 27: 765–783

Jezek J, Chang KT, Joshi AM, Strich R (2019) Mitochondrial translocation of cyclin C stimulates intrinsic apoptosis through Bax recruitment. EMBO Rep 20: e47425

Journo D, Mor A, Abeliovich H (2009) Aup1-mediated regulation of Rtg3 during mitophagy. J Biol Chem 284: 35885–35895

Journo D, Winter G, Abeliovich H (2008) Monitoring autophagy in yeast using FM 4-64 fluorescence. Methods Enzymol 451: 79–88

Kamada Y, Funakoshi T, Shintani T, Nagano K, Ohsumi M, Ohsumi Y (2000) Tor-mediated induction of autophagy via an Apg1 protein kinase complex. J Cell Biol 150: 1507–1513

Kanki T, Wang K, Baba M, Bartholomew CR, Lynch-Day MA, Du Z, Geng J, Mao K, Yang Z, Yen WL et al (2009) A genomic screen for yeast mutants defective in selective mitochondria autophagy. Mol Biol Cell 20: 4730–4738

Kelley LA, Mezulis S, Yates CM, Wass MN, Sternberg MJ (2015) The Phyre2 web portal for protein modeling, prediction and analysis. Nat Protoc 10: 845–858

Khakhina S, Cooper KF, Strich R (2014) Med13p prevents mitochondrial fission and programmed cell death in yeast through nuclear retention of cyclin C. Mol Biol Cell 25: 2807–2816

Klionsky DJ, Codogno P (2013) The mechanism and physiological function of macroautophagy. J Innate Immun 5: 427–433

Lee CW, Wilfling F, Ronchi P, Allegretti M, Mosalaganti S, Jentsch S, Beck M, Pfander B (2020) Selective autophagy degrades nuclear pore complexes. Nat Cell Biol 22: 159–166

Li J, Kim SG, Blenis J (2014) Rapamycin: one drug, many effects. Cell Metab 19: 373–379

Ma M, Burd CG (2020) Retrograde trafficking and plasma membrane recycling pathways of the budding yeast Saccharomyces cerevisiae. Traffic 21: 45–59

Ma M, Burd CG, Chi RJ (2017) Distinct complexes of yeast Snx4 family SNX-BARs mediate retrograde trafficking of Snc1 and Atg27. Traffic 18: 134–144

Ma M, Kumar S, Purushothaman L, Babst M, Ungermann C, Chi RJ, Burd CG (2018) Lipid trafficking by yeast Snx4 family SNX-BAR proteins promotes autophagy and vacuole membrane fusion. Mol Biol Cell 29: 2190–2200

Matscheko N, Mayrhofer P, Rao Y, Beier V, Wollert T (2019) Atg11 tethers Atg9 vesicles to initiate selective autophagy. PLoS Biol 17: e3000377

Millen JI, Krick R, Prick T, Thumm M, Goldfarb DS (2009) Measuring piecemeal microautophagy of the nucleus in Saccharomyces cerevisiae. Autophagy 5: 75–81

Mochida K, Oikawa Y, Kimura Y, Kirisako H, Hirano H, Ohsumi Y, Nakatogawa H (2015) Receptor-mediated selective autophagy degrades the endoplasmic reticulum and the nucleus. Nature 522: 359–362

Mosammaparast N, Pemberton LF (2004) Karyopherins: from nuclear-transport mediators to nuclear-function regulators. Trends Cell Biol 14: 547–556

Murphy R, Wente SR (1996) An RNA-export mediator with an essential nuclear export signal. Nature 383: 357–360

Nagulapalli M, Maji S, Dwivedi N, Dahiya P, Thakur JK (2016) Evolution of disorder in Mediator complex and its functional relevance. Nucleic Acids Res 44: 1591–1612

Nemec AA, Howell LA, Peterson AK, Murray MA, Tomko RJ, Jr. (2017) Autophagic clearance of proteasomes in yeast requires the conserved sorting nexin Snx4. J Biol Chem 292: 21466–21480

Nemet J, Jelicic B, Rubelj I, Sopta M (2014) The two faces of Cdk8, a positive/negative regulator of transcription. Biochimie 97: 22–27

Nice DC, Sato TK, Stromhaug PE, Emr SD, Klionsky DJ (2002) Cooperative binding of the cytoplasm to vacuole targeting pathway proteins, Cvt13 and Cvt20, to phosphatidylinositol 3-phosphate at the pre-autophagosomal structure is required for selective autophagy. J Biol Chem 277: 30198–30207

Papandreou ME, Tavernarakis N (2019) Nucleophagy: from homeostasis to disease. Cell Death Differ 26: 630–639

Popelka H, Damasio A, Hinshaw JE, Klionsky DJ, Ragusa MJ (2017) Structure and function of yeast Atg20, a sorting nexin that facilitates autophagy induction. Proc Natl Acad Sci U S A 114: E10112–E10121

Ramos PC, Hockendorff J, Johnson ES, Varshavsky A, Dohmen RJ (1998) Ump1p is required for proper maturation of the 20S proteasome and becomes its substrate upon completion of the assembly. Cell 92: 489–499

Roberts P, Moshitch-Moshkovitz S, Kvam E, O’Toole E, Winey M, Goldfarb DS (2003) Piecemeal microautophagy of nucleus in Saccharomyces cerevisiae. Mol Biol Cell 14: 129–141

Ronne H, Rothstein R (1988) Mitotic sectored colonies: evidence of heteroduplex DNA formation during direct repeat recombination. Proc Natl Acad Sci U S A 85: 2696–2700

Shintani T, Klionsky DJ (2004) Cargo proteins facilitate the formation of transport vesicles in the cytoplasm to vacuole targeting pathway. J Biol Chem 279: 29889–29894

Shpilka T, Welter E, Borovsky N, Amar N, Shimron F, Peleg Y, Elazar Z (2015) Fatty acid synthase is preferentially degraded by autophagy upon nitrogen starvation in yeast. Proc Natl Acad Sci U S A 112: 1434–1439

Snyder NA, Kim A, Kester L, Gale AN, Studer C, Hoepfner D, Roggo S, Helliwell SB, Cunningham KW (2019) Auxin-Inducible Depletion of the Essentialome Suggests Inhibition of TORC1 by Auxins and Inhibition of Vrg4 by SDZ 90-215, a Natural Antifungal Cyclopeptide. G3 (Bethesda) 9: 829–840

Stanishneva-Konovalova TB, Derkacheva NI, Polevova SV, Sokolova OS (2016) The Role of BAR Domain Proteins in the Regulation of Membrane Dynamics. Acta Naturae 8: 60–69

Stieg DC, Willis SD, Ganesan V, Ong KL, Scuorzo J, Song M, Grose J, Strich R, Cooper KF (2018) A complex molecular switch directs stress-induced cyclin C nuclear release through SCF(Grr1)-mediated degradation of Med13. Mol Biol Cell 29: 363–375

Strich R, Slater MR, Esposito RE (1989) Identification of negative regulatory genes that govern the expression of early meiotic genes in yeast. Proc Natl Acad Sci USA 86: 10018–10022

Suzuki SW, Emr SD (2018) Membrane protein recycling from the vacuole/lysosome membrane. J Cell Biol 217: 1623–1632

Takeshige K, Baba M, Tsuboi S, Noda T, Ohsumi Y (1992) Autophagy in yeast demonstrated with proteinase-deficient mutants and conditions for its induction. J Cell Biol 119: 301–311

Tsai KL, Sato S, Tomomori-Sato C, Conaway RC, Conaway JW, Asturias FJ (2013) A conserved Mediator-CDK8 kinase module association regulates Mediator-RNA polymerase II interaction. Nat Struct Mol Biol 20: 611–619

Uversky VN (2011) Multitude of binding modes attainable by intrinsically disordered proteins: a portrait gallery of disorder-based complexes. Chem Soc Rev 40: 1623–1634

Van Den Hazel HB, Kielland-Brandt MC, Winther JR (1996) Review: biosynthesis and function of yeast vacuolar proteases. Yeast 12: 1–16

Vlahakis A, Lopez Muniozguren N, Powers T (2017) Stress-response transcription factors Msn2 and Msn4 couple TORC2-Ypk1 signaling and mitochondrial respiration to ATG8 gene expression and autophagy. Autophagy 13: 1804–1812

Wang K, Yan R, Cooper KF, Strich R (2015) Cyclin C mediates stress-induced mitochondrial fission and apoptosis. Mol Biol Cell 26: 1030–1043

Wang R, Solomon MJ (2012) Identification of She3 as an SCF(Grr1) substrate in budding yeast. PLoS One 7: e48020

Wanke V, Pedruzzi I, Cameroni E, Dubouloz F, De Virgilio C (2005) Regulation of G0 entry by the Pho80-Pho85 cyclin-CDK complex. EMBO J 24: 4271–4278

Weirich CS, Erzberger JP, Flick JS, Berger JM, Thorner J, Weis K (2006) Activation of the DExD/H-box protein Dbp5 by the nuclear-pore protein Gle1 and its coactivator InsP6 is required for mRNA export. Nat Cell Biol 8: 668–676

Welter E, Thumm M, Krick R (2010) Quantification of nonselective bulk autophagy in S. cerevisiae using Pgk1-GFP. Autophagy 6: 794–797

Willis SD, Hanley SE, Beishke T, Tati PD, Cooper KF (2020) Ubiquitin-proteasome-mediated cyclin C degradation promotes cell survival following nitrogen starvation. Mol Biol Cell 31: 1015–1031

Willis SD, Stieg DC, Ong KL, Shah R, Strich AK, Grose JH, Cooper KF (2018) Snf1 cooperates with the CWI MAPK pathway to mediate the degradation of Med13 following oxidative stress. Microbial Cell 5: 357–370

Yoshida K, Blobel G (2001) The karyopherin Kap142p/Msn5p mediates nuclear import and nuclear export of different cargo proteins. J Cell Biol 152: 729–740

Zhang H, Chen J, Wang Y, Peng L, Dong X, Lu Y, Keating AE, Jiang T (2009) A computationally guided protein-interaction screen uncovers coiled-coil interactions involved in vesicular trafficking. J Mol Biol 392: 228–241

Zhang H, Huang T, Hong Y, Yang W, Zhang X, Luo H, Xu H, Wang X (2018) The Retromer Complex and Sorting Nexins in Neurodegenerative Diseases. Front Aging Neurosci 10: 79

Zhu J, Deng S, Lu P, Bu W, Li T, Yu L, Xie Z (2016) The Ccl1-Kin28 kinase complex regulates autophagy under nitrogen starvation. J Cell Sci 129: 135–144

Zientara-Rytter K, Subramani S (2020) Mechanistic Insights into the Role of Atg11 in Selective Autophagy. J Mol Biol 432: 104–122

Zubenko GS, Park FJ, Jones EW (1983) Mutations in PEP4 locus of Saccharomyces cerevisiae block final step in maturation of two vacuolar hydrolases. Proc Natl Acad Sci U S A 80: 510–514

